# Parvalbumin interneuron inhibition onto anterior insula neurons projecting to the basolateral amygdala orchestrates aversive taste memory retrieval

**DOI:** 10.1101/2021.03.01.433377

**Authors:** Adonis Yiannakas, Sailendrakumar Kolatt Chandran, Haneen Kayyal, Nathaniel Gould, Mohammad Khamaisy, Kobi Rosenblum

## Abstract

Memory retrieval refers to the fundamental ability of organisms to make use of acquired, sometimes inconsistent, information about the world. While memory acquisition has been studied extensively, the neurobiological mechanisms underlying memory retrieval remain largely unknown. The anterior insula (aIC) is indispensable in the ability of mammals to retrieve associative information regarding tastants that have been previously linked with gastric malaise. Here, we show that aversive taste memory retrieval promotes cell-type-specific activation in the aIC. Aversive, but not appetitive taste memory retrieval, relies on specific changes in activity and connectivity at parvalbumin (PV) inhibitory synapses onto aIC pyramidal neurons projecting to the basolateral amygdala. PV aIC interneurons, coordinate aversive taste memory retrieval, and are necessary for its dominance when conflicting internal representations are encountered. This newly described interaction of PV and a subset of excitatory neurons can explain the coherency of aversive memory retrieval, an evolutionary pre-requisite for animal survival.

**Graphical Abstract:** 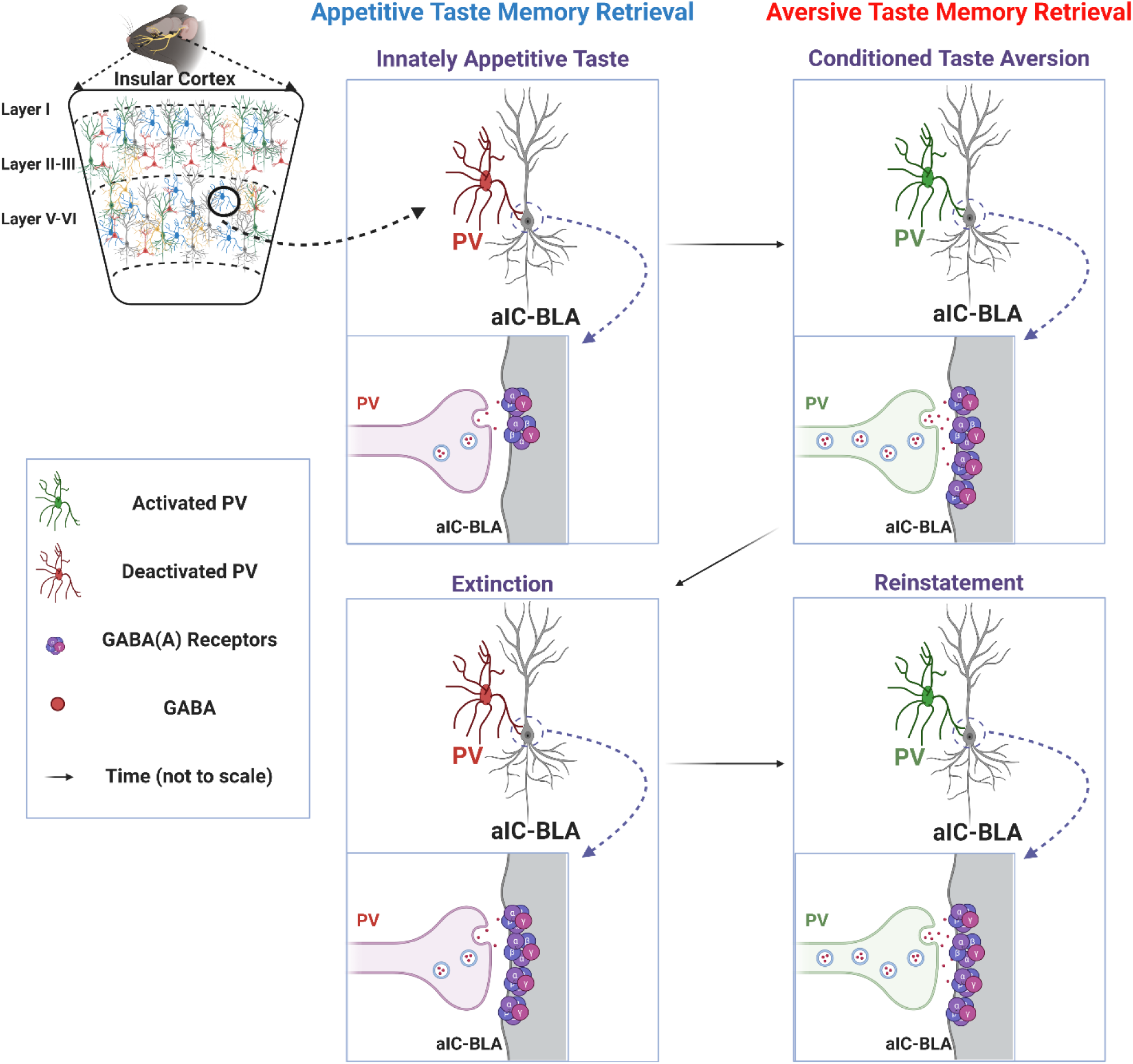

- Retrieval of Conditioned Taste Aversion (CTA) memories at the anterior insular cortex activates Parvalbumin (PV) interneurons and increases synaptic inhibition onto activated pyramidal neurons projecting to the basolateral amygdala (aIC-BLA).
- Unlike innately appetitive taste memory retrieval, CTA retrieval increases the amplitude and frequency of synaptic inhibition onto aIC-BLA projecting neurons, that is dependent on activity in aIC PV interneurons.
- Activation of aIC PV interneurons is necessary for the expression of learned taste avoidance, in both sexes, regardless of stimulus identity.
- Extinction of aversive taste memories suppresses the frequency, but not the amplitude of synaptic inhibition on aIC-BLA projecting neurons.
- The reinstatement of aversive taste memories following extinction is dependent upon activation of aIC PV interneurons and increases in the frequency of inhibition on aIC-BLA projecting neurons.

## Introduction

Taste memories are particularly well suited for the study of learning and memory^1^. In the natural setting, mammals approach novel tastants tentatively, as to assess their associated consequences^2^. Depending on the outcome, a positive or negative memory is formed, that may be retrieved in future encounters, guiding approach or avoidance. Laboratory animals can be trained to express aversion towards innately appetitive tastants (e.g. saccharin, the conditioned stimulus – CS), following a single pairing with malaise inducing agents, such as lithium chloride (LiCl; the unconditioned stimulus - US)^1, 3^. Conditioned taste aversion (CTA) learning results in the formation of an aversive memory that drives CS avoidance in future encounters. This aversive taste memory can be extinguished through unreinforced re-exposure to the CS, and can be reinstated through re-exposure to the original US, among other interventions^4, 5^. The anterior insular cortex (aIC) coordinates taste learning, including CTA, largely independently of the hippocampus^6^. The neurobiological mechanisms of CTA acquisition have been extensively studied ^7, 8^. The processing of avoidance and palatability is further coordinated in concert with other regions, such as the basolateral amygdala (BLA), through changes in activity, connectivity and gene expression^7^. Despite evidence suggesting an indispensable role for the aIC in CTA memory retrieval^9^, research on the underlying cellular and circuit mechanisms is still in its infancy. By using in vivo calcium imaging during CTA retrieval, we have previously demonstrated increased recruitment of aIC-BLA projecting neurons in a valence-, but not stimulus-specific manner^10^, and that activation of this projection is necessary for CTA acquisition, as well as retrieval^11^. More recent evidence would suggest that memory retrieval relies on fine-tuning of activity in excitatory aIC circuits, that is driven by learning-induced changes in connectivity and plasticity^12^. Such learning-induced reconfigurations are thought to be supported by plasticity in inhibitory synapses^13^. For example, during taste memory acquisition in the aIC, GABA_A_ receptors remain initially inactive, however they are activated to inhibit acetylcholine release during retrieval^14^. Earlier studies found that GABA_A_ receptor agonist micro-infusion in the aIC suppresses CTA retrieval^15^, while both agonist and antagonist treatments disrupt appetitive taste memory retrieval^14^. Considering the heterogeneity associated with interneuronal connectivity and activity, little can be concluded regarding the role of GABAergic activity in the aIC during CTA retrieval from existing pharmacological studies^16^. Evidence currently suggests that distinct activity-dependent pathways are recruited across the brain in relation to the local circuitry involved, as well as the aspect of experience being encoded^17^.

The goal of our studies was to identify cellular mechanisms that are indispensable to aversive taste memory retrieval. Utilizing chemogenetic tools in the aIC we show that memory acquisition requires activation of excitatory neurons and inhibition of inhibitory neurons, whereas memory retrieval can be impaired by inhibiting either excitatory or inhibitory neurotransmission (Figure 1). This led us to hypothesize that following CTA learning, the necessity to recruit specific excitatory ensembles for aversive taste retrieval^10, 11^, is accompanied with a necessity for GABAergic activity. By employing electrophysiological recordings from aIC-BLA projecting neurons^10, 11^, we demonstrate that aversive taste memory retrieval indeed promotes valence-specific increases in GABAergic inhibition onto this circuit (Figure 2). This form of inhibitory plasticity is mediated by fast-spiking parvalbumin (PV) aIC interneurons (Figure 3), and is necessary for aversive, but not appetitive taste memory retrieval (Figures 4-5)^16^.

**Figure 1:**
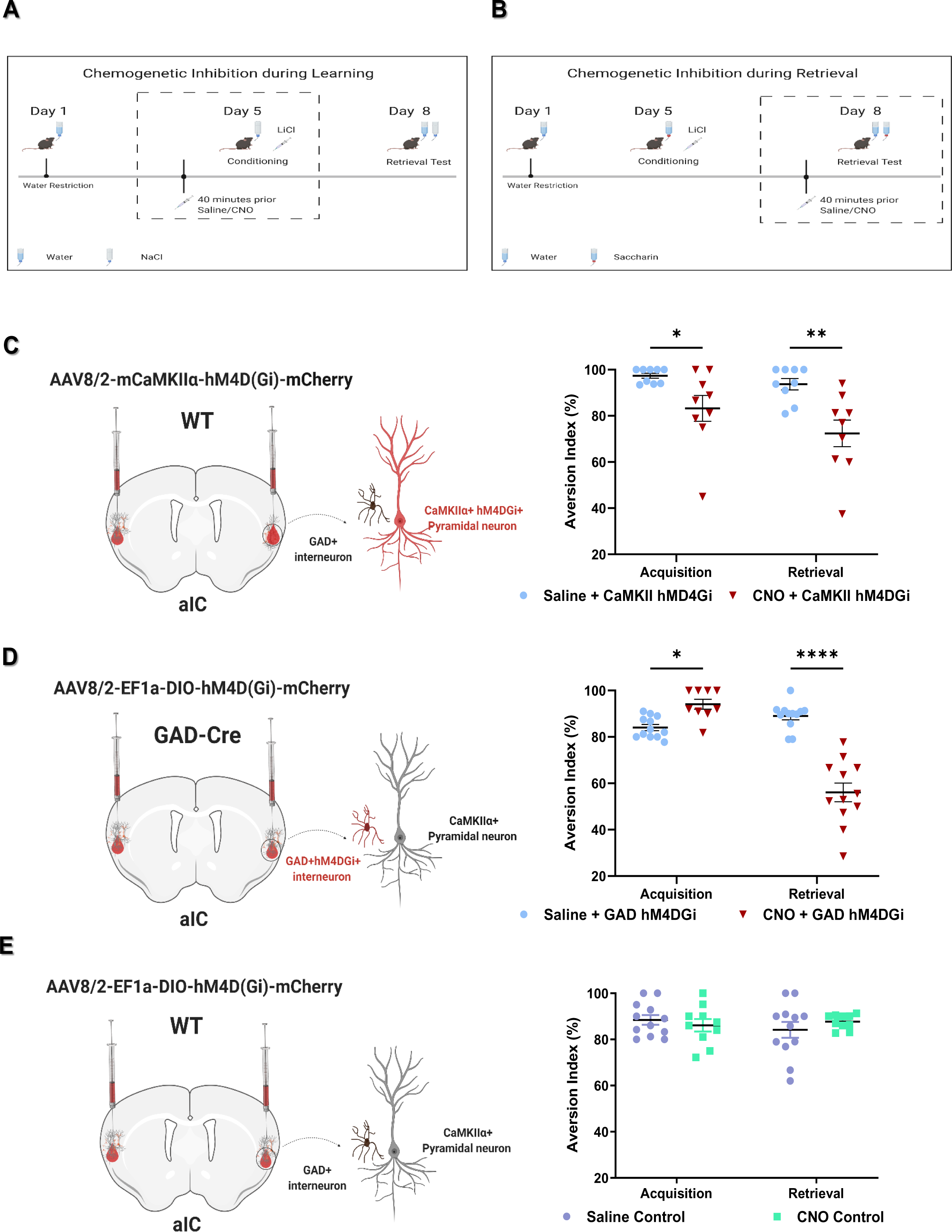
Inhibitory interneurons of the aIC differentially contribute to CTA learning and retrieval. A-B: Cell-type specific chemogenetic silencing of the aIC during CTA acquisition (A) or retrieval (B) was carried out using CNO administration, while control animals received a weight-adjusted dose of saline. C: WT mice were injected with an AAV8 vector resulting in CaMKII-driven expression of hM4DGi-mCherry (left panel). Inhibition of CaMKII aIC neurons during Acquisition suppressed subsequent memory retrieval (Saline – 97.35+/−1.057%, n=9; CNO – 83.23+/−5.605%, n=9). Inhibition of aIC CaMKII during Retrieval, significantly suppressed avoidance (Saline – 93.68+/−2.467%, n=9; CNO - 72.39+/−5.791%, n=9) D: GAD-Cre mice were injected with an AAV8 vector resulting in EF1-driven Cre-dependent expression of hM4DGi-mCherry (left panel). Inhibition of aIC GAD during CTA memory acquisition, enhanced CTA memory retrieval (Saline – 84.04+/−1.315%, n=12; CNO – 94.09+/−2.126%, n=9). Inhibition of GAD aIC interneurons during memory retrieval, significantly suppressed CTA (Saline – 89.00+/− 1.645%, n=12; CNO – 56.08+/−4.026, n=12). E: Control WT mice were injected with an AAV8 vector resulting in Cre-dependent EF1-driven expression of hM4DGi-mCherry (left panel). Administration of a similar dose of CNO in control animals either prior to learning (Saline – 88.46+/−2.090%, n=12; CNO – 86.10+/−2.701%, n=10) or retrieval (Saline – 84.14+/−3.425%, n=12; CNO – 87.71+/−0.9954%, n=10), did not affect avoidance behavior. Avoidance was calculated based on Aversion Index (% +/− SEM). Statistical differences were assessed using 2-way ANOVA (p<0.05(*), p<0.01(**), p<0.001(***), p<0.0001(****)).

**Figure 2:**
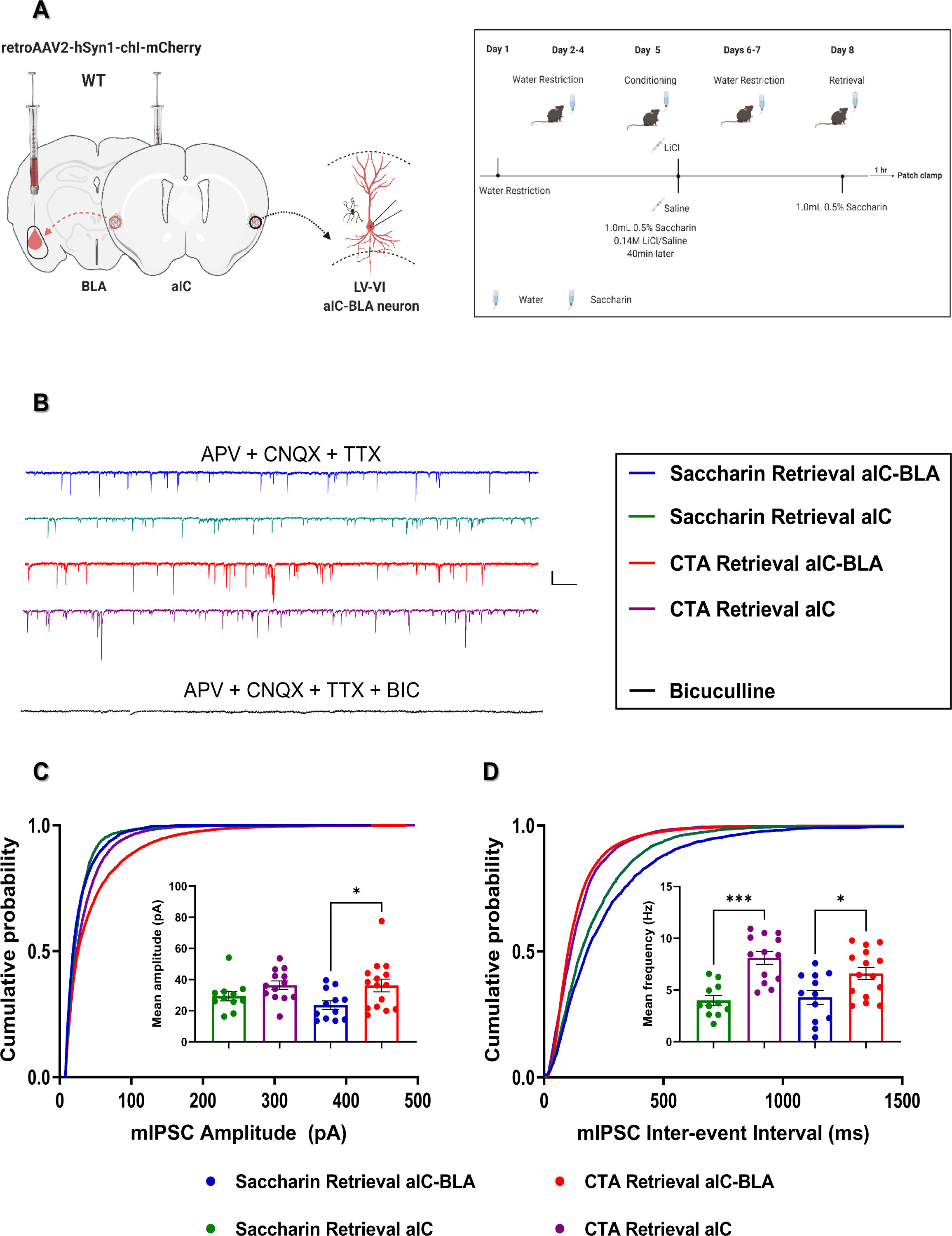
Aversive taste memory retrieval increases mIPSCs on aIC-BLA projecting neurons compared to appetitive memory retrieval. A: WT mice were virally treated to label aIC-BLA projecting neurons using mCherry (n=4-6 mice). B: GABAergic currents on aIC-BLA projecting neurons, or general population of aIC pyramidal neurons (aIC), were isolated in mice retrieving CTA (CTA Retrieval) and stimulus- and familiarity-matched controls (Saccharin Retrieval). Representative traces of mIPSCs recorded from aIC and aIC-BLA-projecting neurons following Saccharin or CTA Retrieval. These events are blocked by Bicuculine. Scale bar represents 50 pA and 500ms. C: Aversive, but not appetitive taste memory retrieval, increased the amplitude of mIPSCs on the BLA projection (Saccharin Retrieval 23.628+/−2.748pA, n=12 cells; CTA Retrieval 36.209+/−4.080pA, n=15 cells). Differences between the two groups in the GP failed to reach significance (Saccharin Retrieval 29.373+/−2.984pA, n=11 cells; CTA Retrieval 36.387+/−2.779pA, n=13 cells). D: The frequency of mIPSCs increased following CTA retrieval compared to controls in both the GP (D – Saccharin Retrieval 3.988+/−0.474Hz, n=11 cells; CTA Retrieval 8.103+/−0.622Hz, n=13 cells) and aIC-BLA projection (D – Saccharin Retrieval 4.292+/−0.6832Hz, n=12 cells; CTA Retrieval 6.596+/− 0.5924Hz, n=15 cells). Statistical analysis was carried out separately for each parameter using 2-way ANOVA (p<0.05(*), p<0.01(**), p<0.001(***), p<0.0001(****)).

**Figure 3:**
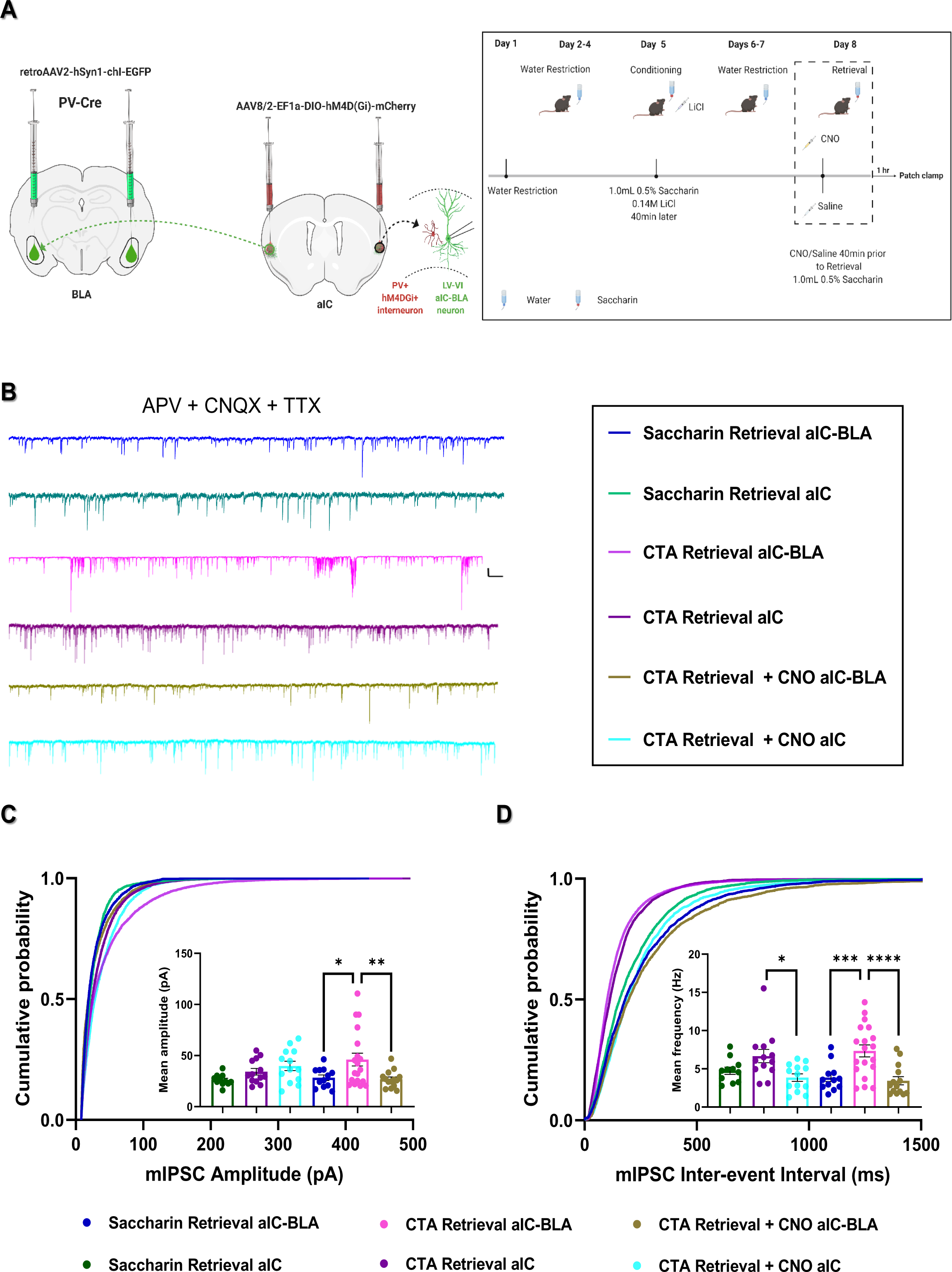
Aversive taste memory retrieval increases PV-dependent mIPSCs on aIC-BLA projecting neurons compared to appetitive memory retrieval. A: Three groups of PV-Cre mice were treated with viral constructs to label aIC-BLA projecting neurons, while inhibitory DREADD receptors were expressed in aIC PVs (n=5-6 mice/group). B: We measured mIPSCs from general population aIC neurons (aIC) and aIC-BLA-projecting neurons in mice following appetitive Saccharin Retrieval or following CTA Retrieval for the same tastant. A third group underwent chemogenetic inhibition of aIC PV during CTA retrieval. Representative traces of mIPSCs recorded from aIC and aIC-BLA-projecting neurons in the 3 treatment groups. Scale bar represents 50 pA and 200ms. C: The amplitude of mIPSCs in the GP was similar among the three treatments (Saccharin Retrieval – 26.153+/−1.663pA, n=11; CTA Retrieval – 34.196+/−3.218pA, n=13; CTA Retrieval + CNO – 39.868+/−4.535pA, n=13). In the aIC-BLA projecting neurons, mIPSC amplitude following CTA Retrieval (46.072+/−6.300pA, n=19) was significantly increased compared to both the Saccharin Retrieval (28.325+/−2.734pA, n=12) and CTA Retrieval + CNO groups (26.797+/−2.251, n=14). D: The frequency of mIPSCs in the aIC following CTA Retrieval (6.663+/−0.877Hz, n=13) was significantly increased compared to CTA Retrieval + CNO (3.854+/−0.489Hz, n=13), but not compared to Saccharin Retrieval (4.750+/−0.473Hz, n=11). mIPSC frequency on aIC-BLA projecting neurons in the CTA Retrieval group (7.341+/−0.778Hz, n=19), was increased compared to both the Saccharin Retrieval (3.856+/− 0.520Hz, n=12) and CTA Retrieval + CNO groups (3.435+/−0.532Hz, n=14). Statistical analysis was carried out separately for each parameter using 2-way ANOVA (p<0.05(*), p<0.01(**), p<0.001(***), p<0.0001(****)).

**Figure 4:**
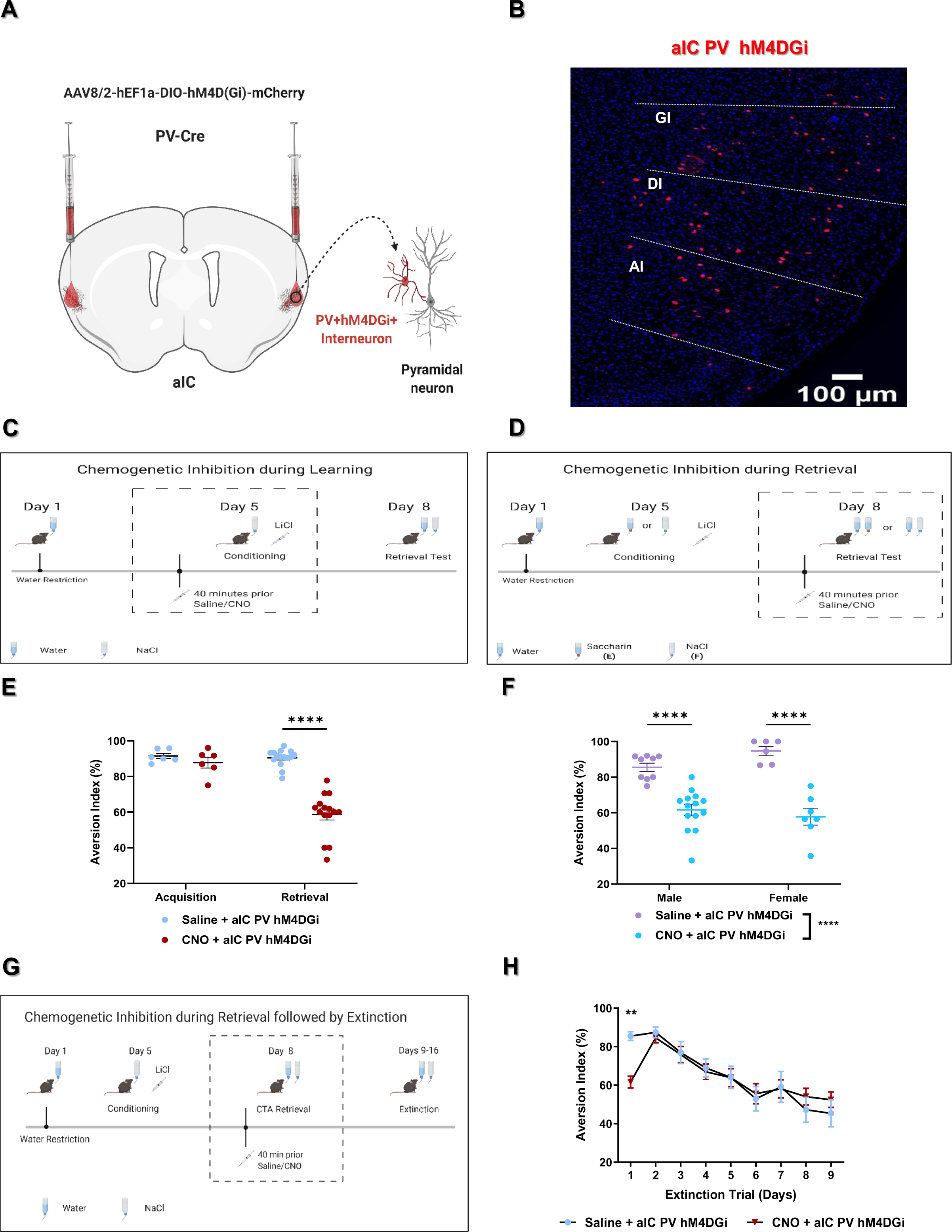
Activation of aIC PV interneurons is necessary for aversive taste memory retrieval, but not for CTA acquisition or maintenance. A-B: Inhibitory DREADDs were introduced in aIC PV interneurons, which was confirmed by immunohistochemistry. Scale bar 100um. C-D: Male and female PV-Cre Mice treated to express inhibitory DREADDs in aIC PV were used to assess their causal role in CTA acquisition and retrieval. E: Chemogenetic silencing of aIC PV in male mice during CTA Acquisition for salty taste did not affect subsequent Retrieval (Saline – 91.47+/−1.453%, n=6; CNO - 87.77+/−3.012%, n=6). Conversely, inhibition of aIC PV during Retrieval significantly suppressed avoidance towards saccharin (Saline – 90.467+/−1.210%, n=15; CNO - 58.701+/−3.141%, n=15). F: Inhibition of aIC PV prior to the Retrieval of CTA for salty taste, significantly suppressed avoidance in both female (Saline – 94.693+/−2.638%, n=6; CNO – 57.78+/−4.685%, n=7) and male mice (Saline – 85.53+/−2.288%, n=9; CNO – 61.01+/−3.079%, n=14). G: Male PV-Cre mice underwent inhibition during CTA retrieval for salty taste, followed by nine unreinforced extinction sessions. H: Inhibition of aIC PV during the retrieval as sufficient to suppress its avoidance (p=0.0065, t==3.436, df=189), but did not subsequently affect the trajectory of extinction (p=0.4885, F (1.189) =0.4818). Statistical differences between the respective groups were assessed using 2-way ANOVA (p<0.05(*), p<0.01(**), p<0.001(***), p<0.0001(****)).

**Figure 5:**
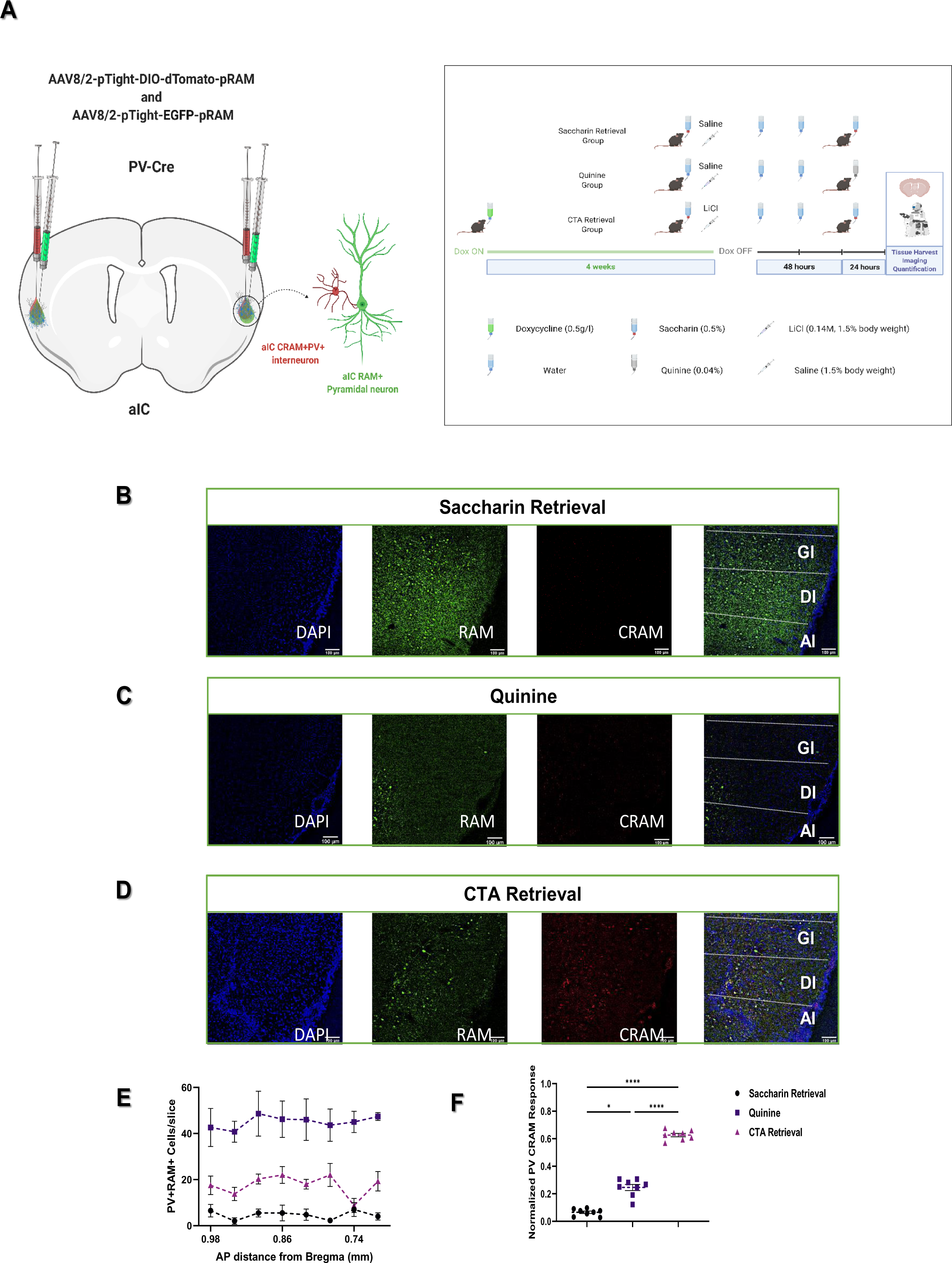
Aversive, but not appetitive taste memory retrieval induces activity-dependent transcription in aIC PV interneurons. A: The CRAM/RAM TET OFF system was utilized in order to label activity-dependent transcription in aIC PV interneurons of PV-Cre mice (see Methods). Activity-dependent transcription was captured in mice following appetitive taste memory retrieval (Saccharin Retrieval, n=4), aversive taste memory retrieval (CTA retrieval, n=5) as well as in response to innately aversive taste exposure (Quinine, n=4). B-D: Representative images from the three treatment groups. Channels displayed from left to right: DAPI, EGFP RAM, TdTomato CRAM, Merge. Scale bars represent 100um. E: Bilateral CRAM cell count data from 8 consecutives slices from the three treatments was plotted in relation to Bregma. Mean PV CRAM response across the aIC were significantly increased in response to CTA Retrieval (45.03+/−0.9114cells/slice) compared to both Quinine (17.69+/−1.591cells/slice), and Saccharin Retrieval (4.688+/− 0.6491cells/slice). Mean aIC PV CRAM responses in the Quinine group were also elevated compared to Saccharin Retrieval. F: Normalized mean CRAM responses among the three treatments. Statistical differences between the groups were assessed using 2-way ANOVA (p<0.05(*), p<0.01(**), p<0.001(***), p<0.0001(****)).

In the natural setting, taste memories are often retrieved along with a host of other information, such that behavior does not merely reflect the choice between gusto and disgust, but rather current subjective perception of the value of the tastant in relation to a spectrum of experiences^2^. In order to distinguish whether aIC PV activation reflects CTA itself, or memory retrieval, we conducted electrophysiological and behavioral studies in mice undergoing CTA extinction and reinstatement. CTA reinstatement following extinction was associated with the re-appearance of PV-dependent modulation of mIPSCs on the aIC-BLA projection, which was necessary and sufficient for aversive memory retrieval (Figure 6). By employing a virally-expressed, cell-type specific activity monitoring system in the aIC^18^, we further show that the recruitment of PV interneurons does not reflect taste identity or familiarity, but rather the dominance of learned aversive memories during retrieval (Figures 5 & 7). Our findings indicate that in the face of conflicting stored information, the ability to retrieve cues that promote the dominance of learned aversive taste memories is dependent on aIC PV activity.

**Figure 6:**
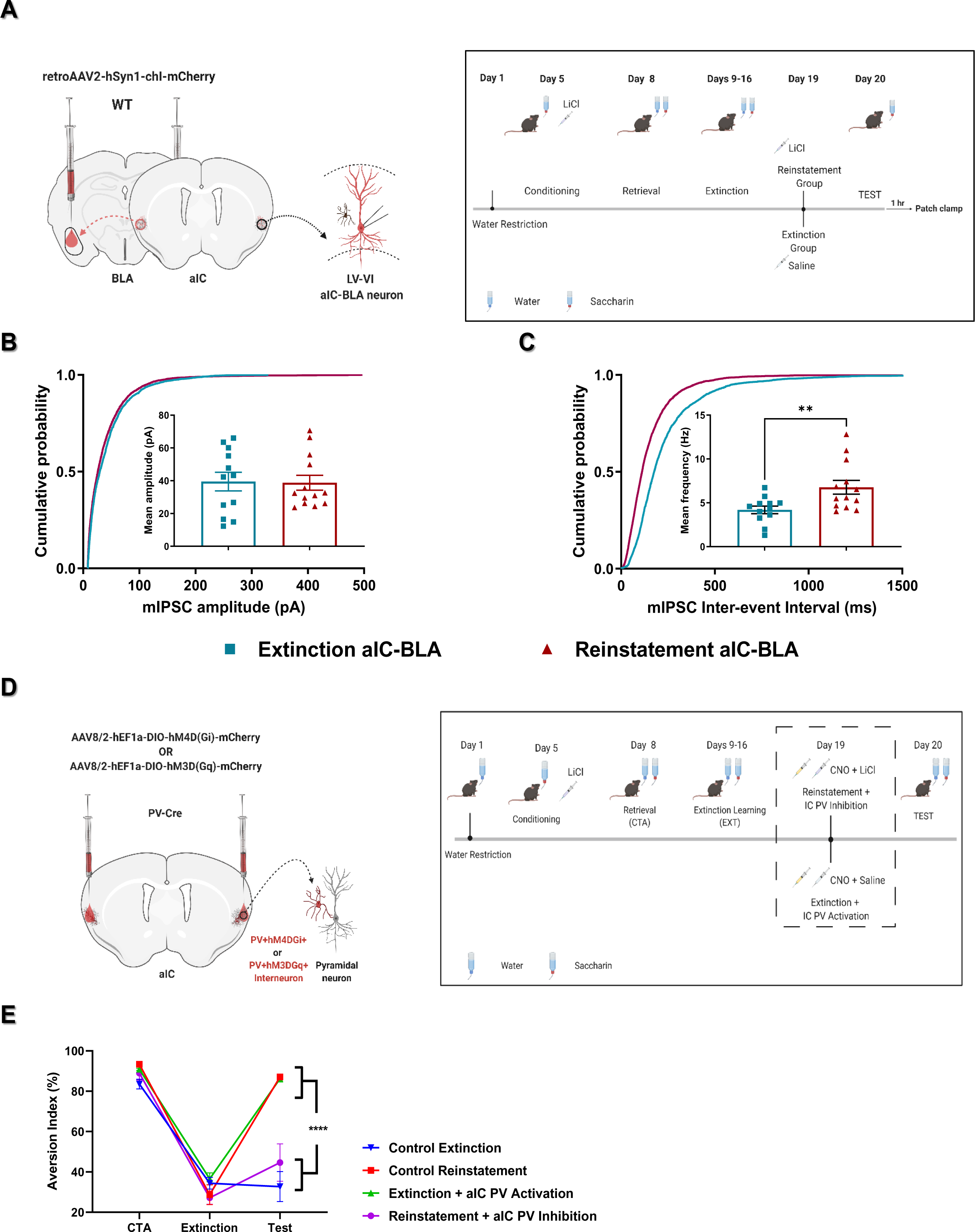
Aversive taste memory retrieval following conflicting experiences is dependent upon mIPSCs driven by aIC PV on aIC-BLA projecting neurons. A: CTA Extinction and Reinstatement studies were conducted in mice treated with viral vectors to label aIC-BLA projecting neurons. Electrophysiological recordings of mIPSCs from LV-VI aIC-BLA projecting neurons (n=3/treatment), were obtained following Extinction and Reinstatement. B: The amplitude of mIPSCs was not different between the two groups (Extinction – 39.49+/−5.606pA, n=12; Reinstatement – 38.76+/−4.519pA, n=13). C: The frequency of mIPSCs was increased in the Reinstatement group, compared to extinction (Extinction – 4.197+/− 0.4367Hz, n=12; Reinstatement – 6.774+/−0.7762Hz, n=13). D: We causally examined the role of aIC PV in reinstatement, using mice expressing activating or inhibitory DREADDs. E: Control (Control Reinstatement; Control Extinction) and treatment groups (Extinction + aIC PV activation; Reinstatement + aIC PV inhibition) exhibited normal CTA retrieval (CTA) and subsequent extinction (EXT). Chemogenetic silencing suppressed CS aversion upon retrieval (Control Reinstatement – 87.088+/−0.732%, n=8; aIC PV inhibition – 37.894+/−9.082%, n=8). Activation 24 hours prior to retrieval reinstated aversive responses following extinction (Control Extinction – 32.702+/−7.434%, n=12; aIC PV activation – 86.24+/−1.613%, n=12). Statistical differences were assessed using Unpaired t-tests (B-C) and 2-way ANOVA (D-E) respectively, (p<0.05(*), p<0.01(**), p<0.001(***), p<0.0001(****)).

## Results

### Inhibitory interneurons of the aIC differentially contribute to CTA learning and retrieval

Neuronal activity within distinct excitatory circuits arising from the aIC differentially contributes to taste memory acquisition and retrieval^11, 19, 20^. We thus hypothesized that activation of excitatory aIC neurons contributes to both CTA memory acquisition and retrieval and assessed this using a chemogenetic viral approach. Briefly, C57Bl6/J wild-type mice (WT) were injected with viral constructs at the aIC, allowing for the incorporation of inhibitory Designer Receptors Exclusively Activated by Designer Drugs (DREADDs) in aIC CaMKII-expressing neurons (Figure 1C; see Methods). In support of our hypothesis, we found that chemogenetic inhibition of aIC CaMKII neurons using Clozapine-N-oxide (CNO) administration prior to CTA acquisition, significantly suppressed CS avoidance upon retrieval testing (Figure 1C, p=0.0496). Similarly, in animals having undergone conditioning without any intervention, inhibition of aIC CaMKII neurons during memory retrieval significantly suppressed aversion (Figure 1C, p=0.0025).

The precise cellular and circuit-wide role of the GABAergic system in aversive taste memory acquisition and retrieval remains unknown, despite significant evidence from pharmacological studies^14, 15^. We hypothesized that, as with excitatory neurons, general activation of GABAergic aIC interneurons is necessary for both CTA learning and retrieval. GAD-Cre mice were injected with viral vectors driving Cre-dependent expression of inhibitory DREADDs in GABAergic aIC interneurons (Figure 1D; Suppl. Figure 1A, C; see Methods). Surprisingly, considering pharmacological studies^14, 15^, inhibition of aIC GAD during CTA acquisition mildly enhanced aversive responses upon retrieval testing (Figure 1D, p=0.0232). Nonetheless, this result would be consistent both with the intricacies in the function among interneuron subtypes^16^, as well as the disinhibition and augmentation of CaMKII neuron function ^21^. Contrariwise, inhibition of aIC GAD acutely suppressed aversive memory retrieval (Figure 1F, p<0.0001). In additional control studies conducted using GAD-CRE WT littermate controls injected with the same virus (Figure 1E), we found that intraperitoneal administration of CNO or saline prior to CTA acquisition (p=0.7686) or retrieval (p=0.5537), does not affect behavioral responses in itself. Even though CTA acquisition favors enhanced excitation in pyramidal aIC neurons^19, 22, 23^, CTA memory retrieval is dependent on activation of GAD interneurons (Figure 1; Suppl. Figure 1), as well as BLA-projecting excitatory neurons^11^. We consequently turned our attention towards examining the possible functional connection between the two neural populations during CTA retrieval.

### Aversive taste memory retrieval increases mIPSCs on aIC-BLA projecting neurons compared to appetitive memory retrieval

Distinct GABAergic interneuron subtypes have been shown to contribute to memory acquisition and consolidation, however little is known regarding their role in memory retrieval^13, 24, 25^. We and others have provided evidence that the output of the aIC to lower brain regions, and the BLA in particular, is instrumental in shaping the valence of learned aversive taste memories^10, 11^. We focused on layer V of the agranular insula, where limbic and gustatory information converge, as well as the majority of aIC-BLA-projecting neurons are localized^10, 11^. Encouraged by the pronounced effects of inhibiting aIC GAD interneurons (Figure 1D), we hypothesized that inhibitory input onto BLA-projecting neurons is induced following aversive, but not appetitive, taste memory retrieval. To test this hypothesis, we measured the frequency and amplitude of miniature inhibitory post-synaptic currents (mIPSCs) from the BLA-projecting population of aIC pyramidal neurons, following taste memory retrieval (Figure 2).

In order to label aIC-BLA-projecting neurons, WT mice were injected with a retroAAV construct in the BLA^11, 26^ (see Methods). Following recovery, one group underwent CTA training for a novel tastant, while a control group underwent familiarization without conditioning (Figure 2A). Electrophysiological recordings of mIPSCs from the general population of pyramidal aIC neurons and aIC-BLA projecting neurons, were then conducted in slices following aversive or appetitive memory retrieval for the same taste (CTA Retrieval or Saccharin Retrieval). To isolate GABAergic currents, we pharmacologically inhibited all excitatory synaptic input and recorded mIPSCs from the projection (Figure 2B).

Consistent with our results using GAD-CRE mice (Figure 1D), aversive, but not appetitive taste memory retrieval was associated with increased amplitude in the BLA projection (Figure 2C, p=0.0179), but not in the general population of aIC pyramidal neurons (Figure 2C, p=0.2901). CTA retrieval also increased the frequency of mIPSCs in the aIC-BLA projection (Figure 2D, p=0.0169), as well as the general population of aIC neurons (Figure 2D, p<0.0001). GABA_A_ receptor antagonism using bicuculline, suppressed both the frequency and amplitude of these events (Figure 2B).

### Aversive taste memory retrieval increases PV-dependent mIPSCs on aIC-BLA projecting neurons compared to appetitive memory retrieval

We next tested the hypothesis that the frequency and amplitude of mIPSCs in the aIC (and the BLA-projecting neurons in particular) observed during CTA retrieval, is dependent upon activation of aIC PV (Figure 3). Cortical PV interneurons have been previously suggested to regulate feedforward inhibition and to shape mIPSCs acting upon excitatory neurons in deep cortical layers^27^. To examine this finding, PV-Cre mice were treated to label aIC-BLA projecting neurons, while inhibitory DREADDs were introduced in aIC PV (Figure 3A-B). Following recovery and CTA acquisition, mice in the treatment group underwent chemogenetic aIC PV inhibition prior to CTA retrieval, while control animals received a saline injection (see Methods).

The frequency and amplitude of mIPSCs recorded from aIC neurons in brain slices following CTA memory retrieval was similar between PV-Cre (Figure 3) and WT mice (Figure 2). In PV-Cre mice, CTA retrieval increased the amplitude and frequency compared to the Saccharin Retrieval group, in aIC-BLA projecting neurons (Figure 3C, p=0.0136; Figure 3D, p=0.0009), but not in the general population of aIC pyramidal neurons (Figure 3C, p=0.5555; Figure 3D, p=0.1854). Inhibition of aIC PV during CTA memory retrieval suppressed both the amplitude (Figure 3C, p=0.0041), as well as the frequency (Figure 3D, p<0.0001) of mIPSCs on aIC-BLA projecting neurons. Importantly, this was not the case in the general population of aIC excitatory neurons, where aIC PV inhibition suppressed the frequency (Figure 3D, p=0.0163) but not the amplitude of mIPSCs (Figure 3C, p=0.7647). Inhibition of aIC PV during retrieval resulted in mIPSCs that closely matched those observed in the appetitive Saccharin Retrieval group across both the aIC, as well as in aIC-BLA projecting neurons (Figure 3C-D). Hence, CTA memory retrieval promotes valence-specific increases in the frequency and amplitude of mIPSCs on aIC-BLA projecting neurons that are at least in part dependent on aIC PV activation. Having identified that PV interneurons are acting upstream to the enhanced inhibition recorded during CTA retrieval at aIC-BLA projecting neurons, we proceeded to examine whether activation of aIC PV is necessary for aversive taste memory retrieval.

### Activation of aIC PV interneurons is necessary for aversive taste memory retrieval, but not for CTA acquisition or maintenance

To address whether the correlative PV-dependent increases in synaptic inhibition at the aIC contribute to CTA memory acquisition or retrieval, PV-Cre mice were treated to express inhibitory DREADDs in aIC PV interneurons (Figure 4; Suppl. Figure 2). Inhibition of aIC PV during CTA training did not affect CS avoidance during retrieval (Figure 4E, p=0.2949). In contrast, aIC PV inhibition prior to the retrieval of CTA for saccharin, significantly suppressed CS avoidance in PV-Cre male mice (Figure 4E, p<0.0001), matching results observed using GAD-Cre mice (Figure 1D).

Using PV-Cre male and female mice retrieving CTA for NaCl, we found that this effect is observed across both sexes, regardless of taste identity (Figure 4F, p<0.0001). Inhibition of aIC PV during CTA retrieval did not affect the subsequent trajectory of extinction in male mice (Figure 4H, 2-way ANOVA, p=0.4885). Post-hoc analysis only identified significant differences during the 1^st^ retrieval session (i.e., CTA memory retrieval), where the treatment was acutely applied (Figure 4H, p=0.0065; see Table of Statistics).

Basic studies have broadly segregated interneurons into three physiologically complementary subtypes, namely parvalbumin (PV), somatostatin (SST), and vasoactive intestinal polypeptide (VIP)-expressing interneurons interneurons^28^. SST interneurons are known to regulate the input to excitatory neurons, by providing powerful dendritic inhibition^28^. VIP interneurons on the other hand, primarily target other interneurons^28^. Complementary experiments in which SST or VIP interneurons were chemogenetically silenced during CTA acquisition or retrieval (Suppl. Figure 1E-F) corroborated that CTA memory retrieval, but not its acquisition, relies on activation of aIC PV interneurons. Our findings encouraged us to test the hypothesis that the recruitment of aIC PVs during CTA retrieval can be defined by monitoring activity-dependent transcriptional programs^18^.

### Aversive, but not appetitive taste memory retrieval increases activity-dependent transcription in aIC PV interneurons

The induction of immediate early genes (IEG) such as Fos, is considered a reliable activity proxy in excitatory neurons, and a number of studies have employed genetically encoded activity reporters to visualize neuronal ensembles activated during memory acquisition and retrieval^29^. The neuron-specific IEG Npas4, has been reported to differ from other markers in its preferential recruitment at inhibitory synapses^30^. Since the focus of our studies was PV activation, we used a viral approach to monitor activity in relation to both *c-fos* and *Npas4* transcription^18^. This AAV construct is available in Cre-dependent (CRAM) and independent variants (RAM), that drive the expression of a DOX OFF system^18^. PV-Cre mice were bilaterally co-injected with CRAM and RAM constructs at the aIC (see Methods). We then allocated the cohort into three experimental groups: one designed to capture aIC PV activity during the retrieval of an appetitive memory for saccharin, a second group retrieving CTA for the same taste, while a third group was exposed to innately aversive quinine (Figure 5; Suppl. Figures 3 & 4). Next, we visualized the spatial distribution of activated aIC neurons (regardless of cell type) through the expression of EGFP (Suppl. Figure 3 - RAM), while activated PVs (Figure 5 - CRAM) were additionally stained with TdTomato (see Methods).

We found the magnitude of RAM response across eight consecutive slices of the aIC was increased in the Saccharin retrieval group (i.e., innately appetitive) compared to both the Quinine (i.e., innately aversive, Suppl. Figure 3E-F, p=0.0043) and CTA saccharin retrieval (i.e., learned aversive, Suppl. Figure 3E-F, p=0.0221) groups. Strikingly, when examining the PV-dependent aIC CRAM response alone (Figure 5), or in relation to the RAM response (Suppl. Figure 4), we found that CTA retrieval increased PV activation compared to both the Saccharin retrieval and Quinine groups (Figure 5E-F, p<0.0001; Suppl. Figure 4B-C, p<0.0001; Suppl. Figure 4D-E, p<0.0001). Importantly, the CRAM response in the Quinine group was significantly increased compared to Saccharin retrieval, where it was hardly detectable (Figure 5E-F, p=0.0102; Suppl. Figure 4D-E, p=0.0002). Our results are consistent with other recent studies, suggesting that suppression of excitatory activity at the aIC during CTA retrieval reflects learning-dependent hedonic shifts in the cortical representation of tastants^12^.

We next sought to examine the ecological and real-life relevance of this newly identified neurobiological mechanism underlying retrieval of one trial aversive experience with an innately appetitive taste. CTA memory retrieval in the laboratory encompasses both CS avoidance, as well as learned aversion towards the tastant itself^2^. However, memory retrieval in the natural setting reflects the balance between a spectrum of positive and negative memories and the availability of cues that promote their reconstruction, as well as relearning and reconsolidation^31^. To address these boundaries we conducted electrophysiological and behavioral studies in mice having undergone CTA extinction and reinstatement^4, 5^, where the expression of avoidance is primarily dependent upon memory dominance and accessibility (Figures 6-7).

**Figure 7:**
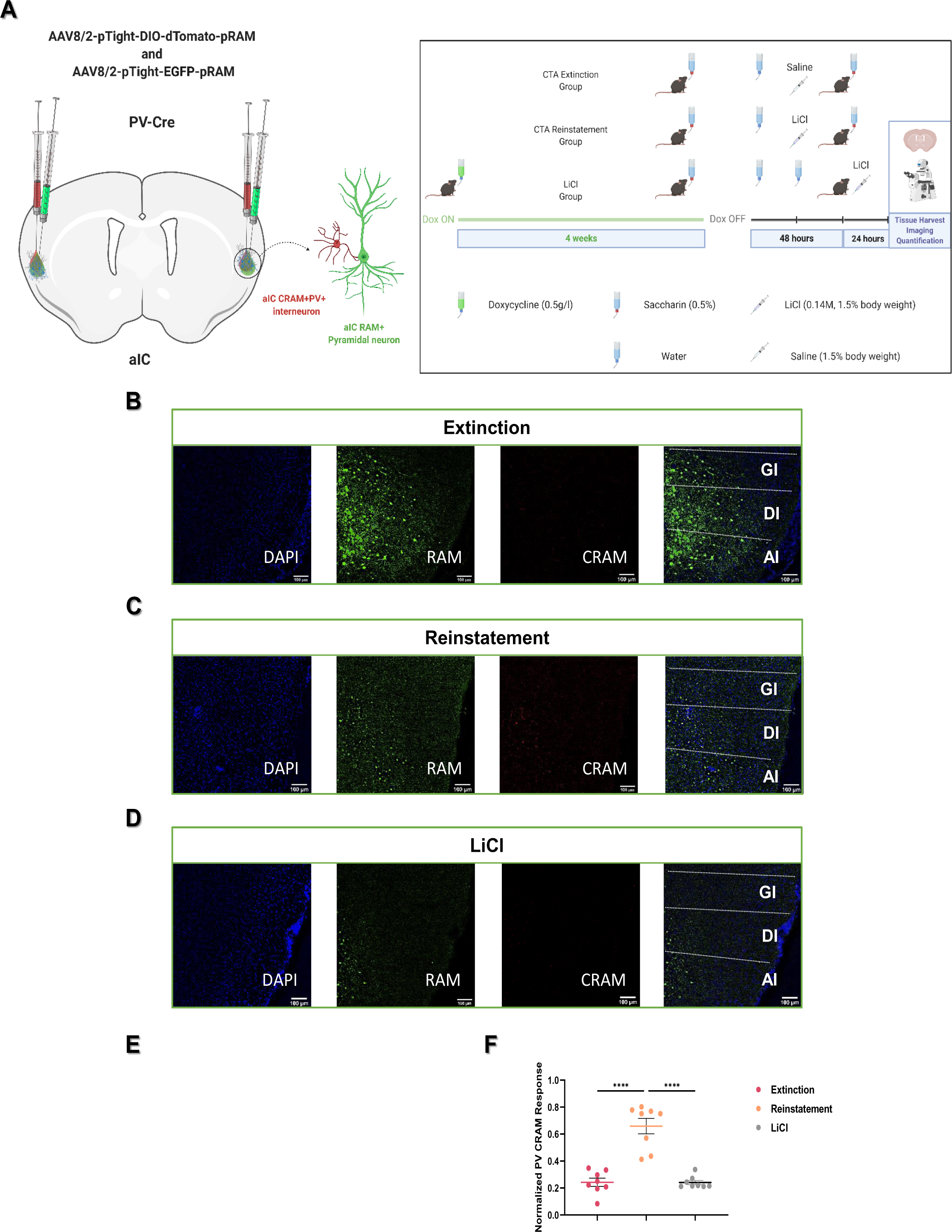
Retrieving aversive taste memory following reinstatement increases activity-dependent transcription in aIC PV. A: The CRAM/RAM TET OFF system was utilized in order to label activity-dependent transcription in aIC PV interneurons of PV-Cre mice (see Methods). Activity-dependent transcription was captured in mice following CTA Extinction (n=3), Reinstatement (n=3), as well as in response to administration of LiCl (n=4). B-D: Representative images from the three treatment groups. Channels displayed from left to right: DAPI, EGFP RAM, TdTomato CRAM, Merge. Scale bars represent 100um. E: Bilateral CRAM cell count data from 8 consecutives slices from the three treatments was plotted in relation to Bregma. Mean PV CRAM response across the aIC were significantly increased in the Reinstatement group (47.38+/−4.104cells/slice) compared to both Extinction (17.42+/−2.181cells/slice) and LiCl (17.31+/−1.122cells/slice). F: Normalized mean CRAM responses among the three treatments. Statistical differences between the groups were assessed using 2-way ANOVA (p<0.05(*), p<0.01(**), p<0.001(***), p<0.0001(****)).

### Aversive taste memory retrieval following conflicting experiences is dependent on PV inhibition of aIC-BLA projecting neurons

To test the hypothesis that aIC PV activation promotes the dominance of aversive memories, we conducted studies of CTA extinction and reinstatement in WT mice (Suppl. Figure 5B). Following conditioning, mice were randomly assigned to the Extinction and Reinstatement groups and underwent a retrieval test (Suppl. Figure 5B; CTA; p=0.8057). Following a total of nine unreinforced CS re-exposures, both groups exhibited preference towards the tastant and extinction of the conditioned response (Suppl. Figure 5B – Extinction; p=0.9967). Animals undergoing US-mediated Reinstatement 48 hours later, exhibited a re-emergence of CS aversion upon testing, unlike the Extinction group (Suppl. Figure 5C – Test, p<0.0001). In a separate cohort of WT mice injected with viral vectors to label aIC-BLA neurons, we recorded mIPSCs from aIC-BLA projecting neurons following CTA Extinction and Reinstatement (Figure 6A-C; Suppl. Figure 5A). The frequency of mIPSCs following reinstatement was significantly increased compared to extinction (Figure 6C, p=0.0021), mirroring CTA retrieval (Figure 2D, 3D). Interestingly, the amplitude of mIPSCs was similar among the two groups (Figure 6B, p=0.0928), and comparable to CTA retrieval (Figures 2C, 3C).

To examine whether the re-emergence of PV-dependent modulation of aIC-BLA projecting neurons causally contribute to reinstatement, we conducted additional chemogenetic studies (Figure 6D-E). Chemogenetic silencing of aIC PV during the administration of the reinstating US, inhibited the re-appearance of aversive behavior upon retrieval testing (Figure 6E, p<0.0001). We further probed our finding in a separate cohort, where we examined whether aIC PV activation following extinction, is sufficient for reinstatement, in the absence of the US. Indeed, chemogenetic activation of aIC PV reinstated CS avoidance (Figure 6E, p<0.0001), proving that aIC PV activation is necessary and sufficient for CTA reinstatement.

### Retrieving aversive taste memory following reinstatement increases activity-dependent transcription in aIC PV

In the final section of our studies, we examined whether the re-emergence of aversive behavior following CTA reinstatement is also associated with cell-type-specific changes in activity-dependent transcription at the aIC (Figure 7; Suppl. Figure 6 & 7). Towards this, we monitored activity in PV-Cre mice using the CRAM/RAM system following Extinction, Reinstatement and the administration of LiCl alone (see Methods). In mice expressing preference towards the CS following extinction, aIC RAM responses closely resembled innately appetitive responses to Saccharin (Suppl. Figure 3). Consequently, aIC RAM in the Extinction group was significantly increased compared to both the Reinstatement (Suppl. Fig 6E-F, p=0.0196) and LiCl groups (p<0.001). Conversely, retrieving reinstated saccharin promoted increased activity-dependent transcription in aIC PVs compared to both the Extinction (Figure 7E-F, p<0.0001) and LiCl groups (p<0.0001). Interestingly, even though LiCl was associated with minimal aIC RAM induction compared to all other groups (see Table of Statistics), a relatively stable portion of this response was contributed by aIC PV (Suppl. Fig 7D-E). Our findings in this section indicate that extinction learning promotes plasticity in CS-evoked internal representations compared to CTA retrieval, as well as the uncoupling of PV interneurons from the activated aIC ensemble. On the other hand, following reinstatement, the dominance of learned aversive memories during retrieval is reflected through the re-emergence of valence-specific circuit configurations reminiscent of CTA retrieval, and the recruitment of PV in the activated aIC ensemble.

## Discussion

The aim of our study was to identify cellular mechanisms subserving coherent memory retrieval and dominance, when encountering conflicting stored information. CTA is an evolutionarily conserved associative learning paradigm, whose acquisition and retrieval are dependent on activity at the aIC^1^. Currently, little is known regarding the biological mechanisms underlying memory retrieval in general, and CTA retrieval specifically^8, 13^. We and others have provided evidence that memory retrieval is improved by interventions that directly or indirectly enhance excitatory input in relevant brain structures during acquisition^21, 22, 25, 32^. Yet, cortical pyramidal neurons exhibit a multitude of activity patterns depending on the behavior and the circuitry involved, while following learning, individual neurons show relatively stable ratios of excitatory and inhibitory conductance^33^. Inhibitory tone is currently thought to impose a synchrony filter onto excitatory activity, that enhances the temporal specificity and dynamic range of firing, as to coordinate behavior in relation to sensory input^16, 34^. Here, we provide evidence that complex cortical operations such as CTA retrieval at the aIC, rely on the entrainment and recruitment of activity in both excitatory and inhibitory circuit components^13, 33^. This learning-dependent entrainment likely takes place over different time scales, that vary depending on the cell types, as well the type of learning involved^35, 36^. Specific excitatory aIC populations recruited during learning can indeed be necessary for memory retrieval (Figure 1). However, our findings point out that the local circuit mechanisms and resultant internal representations necessary for effective memory retrieval, may differ from those engaged in acquisition (Figure 4). CTA memory acquisition initiates a cascade of activity that is not dependent on aIC PV (Figure 4), but likely culminates in consolidation events that prime their necessity for precise memory retrieval^24^. Even so, the necessity for aIC PV activation during CTA memory retrieval does not relate to taste recognition, nor the acquisition of familiarity per se (Figures 5, 7). Instead, activation of aIC PV can be viewed as resultant of a post-learning brain state, that facilitates the accessibility of valence-specific associative information, and memory expression (Figure 6).

The local connectivity of interneuronal networks is highly variable across brain structures, and serves distinct functions in the context of cognition and behavior^37^. The efficacy of inhibitory synaptic transmission, is thought to contribute to homeostatic control and adaptation of neural activity, that enables cortical circuits to exert feed-forward or feedback inhibition in relation to experience^38^. Cortical PV interneurons exhibit promiscuous translaminar connectivity with excitatory afferents, consistent with their role in feed-forward inhibition and gain modulation^39^. Despite their divergent connectivity, cortical PVs require multiple concurrent inputs to spike, tuning their inhibition towards specific pyramidal ensembles^16, 39^. PV interneurons are rapidly activated by afferent inputs, and are swiftly depressed in response to high rates of activity, establishing pockets of network synchrony^40^.

Increases in the amplitude and frequency of mIPSCs, recorded in excitatory cells, are understood to reflect strengthening of the functional connectivity between inhibitory interneurons and pyramidal cells^41, 42^. Such improvements in the efficacy of spontaneous inhibitory synaptic transmission can be achieved by different presynaptic and postsynaptic mechanisms. These include changes in presynaptic GABA (and glycine) vesicular release^43^, changes in the number of postsynaptic receptors^44^, and/or changes in conductance, via modifications in subunit composition or phosphorylation state of postsynaptic receptors^45^. Changes in mIPSC frequency have been attributed to spontaneous release mechanisms recruited at GABAergic presynapses^46^. On the other hand, while increases in mIPSC amplitude may arise through presynaptic means, they are typically the consequence of postsynaptic alterations^44, 45^. We found that aversive, but not appetitive taste memory retrieval over time, relies on increases in the frequency of mIPSCs on the general population of pyramidal aIC neurons, as well as aIC-BLA projecting neurons (Figures 2, 3, 6), that is dependent on PV activity. Importantly, previous studies rule out the possibility of changes in intrinsic properties of infected, BLA-projecting neurons compared to the uninfected general population of aIC neurons ^47^. Our findings would thus indicate that PV interneurons are a major source of inhibitory quantal neurotransmitter release at the aIC, that is necessary for CTA memory retrieval, as to impose a specific firing frequency in both the general, as well as the BLA-projecting pyramidal ensembles (Figures 3 & 4). Further captivatingly, unlike stimulus-matched appetitive controls, mice retrieving CTA exhibited a significant increase in mIPSC amplitude on aIC-BLA projecting neurons, but not in the general population of aIC neurons (Figures 3 & 4). This finding could indicate that the improved synaptic efficacy of aIC-BLA projecting neurons during aversive memory retrieval, is the consequence plasticity on the postsynapse^44, 45^. Future studies are needed to further dissect how such cell-type specific synaptic and intrinsic mechanisms support aversive taste memory retrieval.

In the natural setting, taste memories are often retrieved along with a host of other information, such that behavior does not merely reflect the choice between gusto and disgust, but rather current subjective perception of the value of the tastant, in relation to a spectrum of experiences^2^. As evidenced by our studies in CTA extinction, activation of aIC PV, does not hold the memory itself, and does not contribute to extinction (Figure 4). Instead, the uncoupling of CS-evoked internal representations from the influence of the aIC PV synchrony filter, could be a yet uncharacterized end-point of extinction learning (Figures 6-7). A further interesting perspective is whether the fact that the amplitude of mIPSCs on aIC-BLA projecting neurons is unchanged through the transition from CTA retrieval to extinction and reinstatement (Figure 6-7), is indicative of a “shadow” of the original CTA memory, manifesting on aIC-BLA projecting neurons, that is accessible through PV-dependent modulation^11^. As recent findings elegantly demonstrate, optogenetic activation of aIC PV does not affect chemosensation itself, but can disrupt cognitive signals that drive the execution of taste-guided, reward-directed decision-making^48^. Thus, activation of aIC PV during CTA retrieval likely represents a memory-dependent halt in taste-guided drives towards the CS through the fine-tuning of synaptic activity on aIC-BLA projecting neurons.

Although aIC PV interneurons regulate mIPSCs necessary for aversive memory retrieval, we do not exclude the possibility that other interneuron subtypes contribute either to these mIPSCs themselves, or other phases of learning and memory. Inhibition of PV interneurons, disrupted, but did not abolish mIPSCs from either the general or BLA-projecting portion of the aIC (Figure 3). Indeed, activation of SST can induce reliable inhibition onto PV interneurons, however, PV interneurons themselves provide reciprocal innervation to both SST and VIP interneurons^49^. Further perplexingly, even though VIP preferentially target other interneurons, the majority of inhibition onto PV is mediated by PV-to-PV inhibition and autaptic mechanisms^50^. Sensory experience-dependent breaks in plasticity in the visual cortex, are coordinated through inhibition of SST and VIP interneurons^30^. Such simple correspondence across forebrain structures is often untenable, and further work would be necessary to address how activity within aIC inhibitory circuits contributes to taste learning^13^.

Through the use of the RAM/CRAM activity monitoring system we demonstrate that internal representations recruited at the aIC remain plastic in relation to past knowledge, and obey topography rules that reflect their perceived valence. Consistent with recent studies^12^, we found that the size of the recruited ensemble at the aIC is restricted in response to CTA taste memory retrieval or reinstatement (Figures 5 & 7). Conversely, aIC RAM responses were significantly enhanced in mice expressing preference for the same taste, regardless of familiarity or prior experience (Suppl. Figures 3 & 6). The processing of innately aversive stimuli such as LiCl and quinine, has been shown to be primarily focalized in more caudal portions of the IC^20, 51^. In accord, innately aversive stimuli did not induce significant RAM responses at the aIC (Figures 5 & 7). CRAM labelling in aIC PV was significantly enhanced in response to the retrieval of learned aversion, however labeling was also detectable in response to innately aversive stimuli (Suppl. Figure 4 & 7). This finding could be further consistent with earlier work suggesting that learned aversive responses rely on the hi-jacking innate mechanisms, as to override chemosensory-driven behavior^52^. Studies have argued that the IC is responsive to taste identity, and that sweet- and bitter-responsive cortical fields project to topographically distinct areas of the amygdala^53^. However, the IC is an elongated cortical structure, and our studies did not extend to these rostral, or dorsal extremities of the region (see Methods). Nonetheless the aforementioned findings might not be mutually exclusive, and chemosensation might be differentially distributed within multimodal cortical structures^54^.

Taken together, our results point towards a learning-dependent plasticity mechanism that entrains the GABAergic system in relation to experience, as to impose a temporal code of firing onto specific excitatory circuits during recall. In line with other recent reports, we demonstrate that the ability of distinct brain regions to acquire and retrieve of information, might in fact be subserved by distinct components of the local circuit and over different time scales^21, 32^. Several upstream circuits might be involved in the modulation aIC PV activity during CTA retrieval, including thalamocortical and amygdalocortical projections^55^, as well as the parabrachial nucleus^14^ and basal forebrain^56^. Recent evidence from brain-wide connectivity studies of the mouse insular cortex, corroborate that the portion of the IC examined in our studies is indeed home to local PV interneurons, and is bi-directionally connected with the thalamus and amygdala through excitatory projections^57^. Alternatively, activity-dependent plasticity in aIC-BLA projecting neurons itself^10, 11^, might guide the entrainment and subsequent recruitment of PV interneurons during retrieval^21, 24, 30^.

Even though the primary focus of our studies was the role of aIC PV in taste memory retrieval, we do not presume their role is limited to this function. The IC integrates information in relation to a number of modalities, that subserve social behaviors, drug seeking, as well as metabolic and interoceptive information^58^. Aberrations in memory retrieval represent a common theme among a number of neuropsychiatric disorders, including schizophrenia and autism. Impairments in cognition observed in patients and animal models, have been linked to dysfunction of cortical PV interneurons^34, 59^. Clinical imaging studies have implicated aIC connectivity in the incidence and manifestation of neuropsychiatric disease, however is the methods used in these studies do not allow for the interrogation of the influence of dysfunction in aIC inhibitory circuits^60, 61^. Future research will attempt to dissect the contribution of distinct local and brain-wide circuits arising at the aIC in learning and memory processes. Tools that allow the dissection of multiple activity-regulated cellular components within the same brain structure could be revealing of cell-type specific mechanisms that enable memory formation or retrieval in health and disease.

## STAR Methods

### Animals

Wild type (C57Bl6/J; Stock Number 000664; WT), GAD-Cre and their WT littermates (Gad2tm2(cre)Zjh/J; Stock Number 010802), SST-CRE (SSTtm2.1(cre)Zjh/J; Stock Number 013044), PV-Cre (B6.129P2-Pvalbtm1(cre)Arbr/J; Stock Number 017320), or VIP-Cre (Vip-tm1(cre)Zjh/J; Stock Number 010908) adult male or female mice (8-12 weeks) were used as described. All mice were housed in the local Animal Resource Unit with water and standard chow pellets available ad libitum, under a 12-hour dark/light cycle, except where stated. All procedures were approved by the institutional animal care and use committee in accordance with the University of Haifa regulations and National Institutes of Health Guidelines, under Ethical Licenses 487/17, 505/17 and 636/19.

### Surgery and viral injection

Mice were treated with norocarp (0.5mg/kg), 30 minutes prior to being briefly anesthetized using isoflurane (4-5%) in a suitable chamber (M3000® NBT Israel®/Scivena Scientific®). Following the induction of deep anesthesia, animals were transferred to a Model 963 Kopf® stereotaxic injection system, and maintained through a nose cone (1-2%), for at least 10 minutes. Upon confirming the lack of pain responses, the skull was surgically exposed and following relevant alignment and calibration, mice were injected with appropriate AAV constructs at the anterior agranular insula (AP +0.86; ML ±3.4; DV -4.00) and/or the basolateral amygdala (AP -1.6; ML ±3.375; DV -4.80) as specified in each experiment (see Viral Vectors)^26^.

Viral delivery was performed using a Hamilton micro-syringe (0.1ul/minute). Appropriate cleaning and suturing (Vetbond®) of the exposed skull was performed at the end of viral delivery. Animals were then administered an additional dose of 0.5mg/kg norocarp, as well as 0.5mg/kg of Baytril ® (enrofloxacin), and then transferred to an appropriately clean and heat-adjusted enclosure for 2 hours. Upon inspection, mice were returned to fresh cages along with similarly treated cage-mates. Weight-adjusted doses of the Norocarp and Baytril were administered for an additional 3 days. All AAV constructs used in this study were obtained from the Viral Vector Facility of the University of Zurich (http://www.vvf.uzh.ch). All mice used in our studies were bilaterally injected with AAV constructs, as defined below.

### Table of Viral Vectors

| **Viral Vectors** |  |
| --- | --- |
| **Figure** | **Treatment** |
| **Figure 1C** | WT C57Bl6/J mice were bilaterally injected with AAV8/2-mCamKIIα-hM4D(Gi)_mCherry-WPRE-hGHp(A) at the aIC (0.25uL/site, Physical Titer 3.1 x 10E12 vg/mL) |
| **Figure 1D** | GAD-Cre mice were bilaterally injected with AAV8/2-hEF1a-dlox-hM4D(Gi)_mCherry(rev)-dlox-WPRE-hGHp(A) at the aIC (0.25uL/site, Physical Titer 4.5 x 10E12 vg/mL) |
| **Figure 1E** | GAD-CRE<sup>-</sup> WT littermates were bilaterally injected with AAV8/2-hEF1a-dlox-mCherry(rev)-dlox-WPRE-hGHp(A) at the aIC (0.25uL/site, Physical Titer 4.5 x 10E12 vg/mL) |
| **Figure 2** | WT C57Bl6/J mice were bilaterally injected with retroAAV2-hSyn1-chI-mCherry-WPRE-SV40p(A) at the BLA (0.25uL/site, Physical Titer 6.6 x 10E12 vg/mL) |
| **Figure 3** | PV-Cre mice were bilaterally injected with AAV8/2-hEF1a-dlox-hM4D(Gi)_mCherry(rev)-dlox-WPRE-hGHp(A) at the aIC (0.25uL/site, Physical Titer 4.5 x 10E12 vg/mL), and retroAAV2-hSyn1-chI-EGFP-WPRE-SV40p(A) at the BLA (0.25uL/site, Physical Titer 6.6 x 10E12 vg/mL) |
| **Figure 4, Suppl. Figure 2** | PV-Cre male or female mice were bilaterally injected with AAV8/2-hEF1a-dlox-hM4D(Gi)_mCherry(rev)-dlox-WPRE-hGHp(A) at the aIC (0.25uL/site, Physical Titer 4.5 x 10E12 vg/mL) |
| **Figure 5 & 7** | PV-Cre mice were bilaterally co-injected with AAV8/2-pTight-dlox-dTomato(rev)-dlox-WPRE-bGHp(A)-rp(A)-pRAM-d2tTA-SV40p(A) and AAV8/2-pTight-EGFP-WPRE-bGHp(A)-pRAM-d2tTA-SV40p(A) at the aIC (0.35uL/site, Physical Titer 4.5 x 10E12 vg/mL) |
| **Figure 6A-C, Suppl. Figure 5A** | WT mice were bilaterally injected with retroAAV2-hSyn1-chI-EGFP-WPRE-SV40p(A) at the BLA (0.25uL/site, Physical Titer 6.6 x 10E12 vg/mL) |
| **Figure 6D-E, Suppl. Figure 2** | PV-Cre mice were bilaterally injected with AAV8/2-hEF1a-dlox-hM4D(Gi)_mCherry(rev)-dlox-WPRE-hGHp(A), while a separate cohort was injected with AAV8/2-hEF1a-dlox-hM3D(Gq)_mCherry(rev)-dlox-WPRE-hGHp(A) at the aIC (0.25uL/site, Physical Titer 2.8 x 10E12 vg/mL) |
| **Suppl. Figure 1C, E** | SST mice were bilaterally injected with AAV8/2-hEF1a-dlox-hM4D(Gi)_mCherry(rev)-dlox-WPRE-hGHp(A) at the aIC (0.25uL/site, Physical Titer 4.5 x 10E12 vg/mL) |
| **Suppl. Figure 1D, F** | VIP-Cre mice were bilaterally injected with AAV8/2-hEF1a-dlox-hM4D(Gi)_mCherry(rev)-dlox-WPRE-hGHp(A) at the aIC (0.25uL/site, Physical Titer 4.5 x 10E12 vg/mL). |

### Behavioral procedures

All animals undergoing stereotaxic surgery were allowed 3 weeks of recovery in their home cages. During the 4^th^ week of recovery, mice were transferred to individual cages with ad libitum access to food and water. On the last day of the 4th week of recovery, mice were water-deprived for 24 hours. For the subsequent 48 hours, mice were allowed access to tap water in sterile plastic pipettes for 8 hours/day. Regardless of subsequent behavioral procedures (described below), mice were then water-restricted for an additional 2 days, being allowed access to water for 1 hour, during the early hours of the light cycle.

### Behavioral Studies in Conditioned Taste Aversion Acquisition and Retrieval

On the 5^th^ day from the start of water restriction training, mice were allowed access to 1.0mL of 0.5% saccharin- or 0.5% NaCl-water for 20 minutes^19^. Forty minutes following the start of the 20-minute drinking session (inter-stimulus interval, ISI=40minutes), animals were intraperitoneally injected with a 1.5% body weight dose of the malaise inducing agent LiCl (0.14M), the unconditioned stimulus (US).

Mice undergoing cell-type specific chemogenetic inhibition of the aIC during conditioned taste aversion (CTA) acquisition (Figure 1C, D, E; Figure 4C; Suppl. Figure 1E-F), received CNO (1.0mg/kg - Enzo®) 40 minutes prior to the start of the training session, while control animals received a weight-adjusted dose of saline (1% body weight). Animals undergoing chemogenetic inhibition prior to Acquisition were trained for salty-water (0.5% NaCl). For the subsequent 2 days, mice were allowed access to water for 1 hour, 3 hours from the start of the light cycle. Three days following CTA acquisition, mice were provided with a choice between the conditioned stimulus (CS) and tap water in pipettes.

In experiments designed to examine the effect of cell-type specific chemogenetic inhibition of the aIC during CTA retrieval, male mice underwent CTA training for 0.5% saccharin, without any further interventions (Figure 1C, D, E; Figure 3B-C. Figure 4; Suppl. Figure 1E-F). Three days later, and 40 minutes prior to the start of the retrieval test, mice received 1.0mg/kg of CNO (Enzo®), or 1% body weight saline.

Aversion to the conditioned tastant (0.5% NaCl or 0.5% saccharin) was calculated by expressing the volume of water consumed during the retrieval session, as a percentage of the total intake (Volume of water consumed) / (Volume of water + Volume of tastant consumed)). All data presented as a percentage +/− the standard error of mean (SEM).

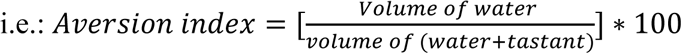

A separate cohort of female PV-Cre mice treated to express inhibitory chemogenetic receptors in aIC PV underwent CTA training for 0.5% NaCl (Figure 4F, H). The effect of aIC PV inhibition during CTA retrieval was examined as above. Additional control studies were conducted in control mice (not expressing CRE), where the same viruses were injected in the aIC (Figure 1E). Following surgery recovery as above, mice in these control groups received 1.0mg/kg CNO or saline (1% body weight) 40 minutes prior to CTA acquisition for 0.5% NaCl, or the retrieval of CTA for 0.5% saccharin.

### Behavioral Studies in CTA Extinction and Reinstatement

WT C57Bl/6J adult male mice used to study extinction and reinstatement were trained in CTA for saccharin (Suppl. Figure 5B). For the subsequent nine drinking sessions, all mice were allowed an unreinforced choice test between the CS and water for 30 minutes. Following these extinction sessions, mice were water-restricted for another 48 hours. The cohort (n=16) was split into the extinction and reinstatement groups (n=8 each). The reinstatement group received an identical intraperitoneal dose to the original US, 24 hours prior to retrieval testing^4, 5^. Conversely, the extinction group received a similarly weight-adjusted dose of saline. Behavior was recorded and analyzed during CTA retrieval (1^st^ session), at the peak of extinction (9^th^ session), and during the final test day (10^th^ session).

Two separate sets of experiments were conducted to address the role of aIC PV interneurons in extinction and reinstatement (Figure 4E and Figure 6). PV-Cre mice were treated to express chemogenetic inhibitory receptors in PV interneurons (Figure 4F) and underwent CTA acquisition for 0.5% NaCl. Mice were then randomly split into the treatment and control groups, receiving CNO (1.0mg/kg) or saline prior to the first retrieval session (Figure 4H). For the subsequent nine days, mice underwent extinction training as above.

PV-Cre mice were treated to express chemogenetic inhibitory and excitatory receptors in PV interneurons of the aIC and examine their causal role in CTA reinstatement (Figure 6D-E). Following recovery and CTA acquisition for 0.5% saccharin, mice were randomly split into the treatment and control groups. Both groups underwent CTA extinction, as described. A portion of the mice expressing inhibitory DREADDs, received 1.0mg/kg of CNO 40 minutes prior US-mediated reinstatement, while mice expressing excitatory DREADDs received CNO followed by a saline injection using the same time framework. Control mice in the inhibitor group received saline followed by the US, while control mice expressing excitatory DREADDs received two saline injections within the same timeframe (1% body weight). Behavioral responses were recorded 24 hours later (Figure 6D-E).

### Electrophysiological Studies of CTA Retrieval

WT C57Bl6/J mice were used to examine the synaptic properties of aIC-BLA projecting neurons and the general population of aIC pyramidal neurons (Figure 2). Mice were treated with viral constructs labeling aIC-BLA projecting neurons (see Viral Vectors). Upon recovery, mice in CTA retrieval group were trained in CTA for saccharin, while the Saccharin Retrieval group received a matching body weight-adjusted injection of saline. Three days following conditioning, both groups underwent a memory retrieval task, receiving 1.0mL of the CS 1 hour prior to sacrifice under deep anesthesia. Brain tissue was extracted and prepared for electrophysiological recordings (described below).

PV-Cre mice were used to examine the influence of aIC PV activity in the incidence of mIPSCs on aIC-BLA projecting neurons (Figure 3, see Viral Vectors). Following recovery and water restriction, mice were randomly split into three groups. The Saccharin Retrieval group underwent familiarization with the tastant without conditioning as above. The CTA Retrieval and CTA retrieval + CNO groups were trained in CTA for saccharin, and then randomly split into their assigned groups. The CTA retrieval + CNO group received CNO (1.0mg/kg) 40 minutes prior to a retrieval session where they were allowed 1.0mL of the CS. The CTA Retrieval group received a 1% body weight dose of saline at the same time point. All groups were sacrificed under deep anesthesia, 1 hour from the start of the retrieval session. Brain tissue was prepared for subsequent recordings.

### Electrophysiological Studies of CTA Extinction and Reinstatement

CTA extinction and reinstatement electrophysiological studies were conducted in a cohort of WT male mice (Figure 6A-C). Following surgery, recovery and water restriction training, animals were randomly assigned to the extinction and reinstatement groups. PV-Cre adult male mice used to study extinction and reinstatement were trained in CTA for saccharin Following extinction, the reinstatement group received an identical intraperitoneal dose to the original US, 24 hours prior to retrieval^4, 5^. Conversely, the extinction group received a similarly weight-adjusted dose of saline. During the final retrieval session, both groups of mice were allowed access to 1.0mL of the CS, 1 hour prior to sacrifice under deep anesthesia and slice preparation for electrophysiology.

### Electrophysiology

#### Insula slice preparations

Following behavioral procedures as described for Figures 2 and 4, mice were deeply anesthetized with 5% isoflurane and transcardially perfused with 40mL of ice-cold oxygenated cutting solution containing (in mM): 25 NaHCO_3_, 105 Choline-Chloride, 2.5 KCl, 7 MgCl_2_, 0.5 CaCl_2_, 1.25 NaH_2_PO_4_, 25 D-Glucose, 1 Na-Ascorbate and 3 Na-Pyruvate. 300 µm thick coronal brain slices were cut with a Campden-1000 Vibrotome using the same cutting solution starting approximately at Bregma 1.34. Slices were incubated for at least 60 minutes at 34°C in artificial cerebrospinal fluid (ACSF) containing (in mM): 125 NaCl, 25 NaHCO_3_, 2.5 KCl, 1.25 NaH_2_PO_4_, 2 CaCl_2_, 1 MgCl_2_ and 25 D-Glucose, prior to transfer to the electrophysiological setup. During electrophysiological recordings, slices were placed in an ACSF-perfused recording chamber (2.0mL/min, 32-34°C). All solutions were constantly carbogenated with carbogen (95% O2 + 5% CO_2_).

#### Whole-cell recording

Pyramidal cells were visualized using infrared light illumination, under differential interference contrast microscope (DIC) with 40x, water-immersion objective mounted on a fixed-stage microscope (BX51-WI; Olympus). aIC-BLA-projecting neurons were identified by the presence of mCherry (Figure 2) or EGFP (Figure 3). The expression of inhibitory DREADD receptors in PV interneurons of the IC was confirmed by the identification of mCherry-positive cells (Figure 3). Whole cell recordings from aIC-BLA-projecting neurons and neurons from the general aIC pyramidal population (no fluorescence), were performed using an Axopatch 200B amplifier and digitized by Digidata 1440 (Molecular Devices). Recording electrodes were pulled from a borosilicate glass pipette (3–5 MΩ) using an electrode puller (P-1000; Sutter Instruments). Recordings were made from the soma of insula LV-VI pyramidal cells. Voltages for liquid junction potential (+10 mV) were not corrected online. Series resistance was compensated and only series resistance <20 MΩ was included in the data set. Pipette capacitance was ∼80% compensated. To record GABA_A_-receptor-mediated miniature IPSCs (mIPSCs), the internal solution contained (in mM): 140 cesium chloride, 1 EGTA, 6 KCl, 4 NaCl, 2 MgCl2, and 10 HEPES, pH 7.25 and 280 mOsm. In these conditions, the reversal potential of chloride is ∼0 mV. (CNQX) 20 µM (Sigma), (D-AP-5) 50µM (Tocris) and tetrodotoxin (TTX) 1 µM were also added to the external ACSF solution. 50-µM bicuculline (Tocris) were added to the bath to block GABA_A_ receptors. Data were acquired with pClamp 10.7 (Molecular Devices) and recorded with an Axopatch 200B amplifier (Molecular Devices). Data sampled at 20 kHz and filtered at 2 kHz. Series resistance, Rin, and membrane capacitance were monitored throughout, and experiments where resistance changed >20%, were discarded. mIPSCs were recorded in voltage clamp mode at a holding potential −60 mV. Data analysis was carried out using Clampfit (Molecular Devices, Sunnyvale, CA).

### Immunohistochemistry

#### Tissue preparation

Brain tissue from all behavioral experiments was collected under deep anesthesia, following transcardial reperfusion using 4% paraformaldehyde solution (PFA). Tissue was incubated for 24 hours in PFA, followed by 48 hours in 30% sucrose in phosphate buffered saline (PBS). Tissue was subsequently frozen at −80°C and processed for slicing using a cryostat (Leica CM 1950®). Eight 35μm-thick brain slices were collected between Bregma 0.98 and 0.70, and treated for fluorescent immunohistochemistry. Immediately following slicing, tissue was briefly washed in PBS, prior to blocking and permeabilization for 1 hour using a 0.3% bovine serum albumin/0.3% Triton X-100/10% fetal bovine serum solution in PBS (blocking solution). Slices were washed and mounted on glass slides using Vectashield® Mounting Medium (H-1000). Slides were visualized using a vertical light microscope (Olympus CellSens Dimension ®) at 10x and 20x magnification.

#### Activity-dependent labelling using viral expression of the RAM system at the aIC

Adult male PV-Cre mice were co-injected with viral constructs resulting in CRAM and RAM vectors (Figure 5 & 7, Suppl. Figures 3-4, Suppl. Figures 6-7). Following 3 days of recovery, mice were provided with doxycycline supplemented water (0.5g/L) for 4 weeks. During the 4^th^ week mice were split into individual cages and were trained to drink doxycycline-supplemented water from pipettes, through two daily 1 hour sessions, at the start and end of the light cycle^18^. Mice were then randomly assigned to respective groups. As to open the window for RAM-dependent labeling^18^, all mice were allowed 48 hours of two 1-hour water-drinking sessions/day prior to stimulus presentation. Mice were sacrificed twenty-four hours following stimulus presentation^18^.

##### Saccharin Retrieval Group

On the last day of doxycycline supplementation, mice received 1.0mL of 0.5% saccharin followed by an intraperitoneal injection of saline (1.5% body-weight). They were then allowed access to 4.0mL of doxycycline-water. Three days later, the mice were exposed to 1.0mL of the tastant for the 2^nd^ time. Twenty-four hours later, mice were sacrificed following a choice test between water and the tastant.

##### Quinine Group

The Quinine group was similarly maintained on doxycycline, but was not exposed to saccharin. On the last day of the 4^th^ week on doxycycline, mice in the Quinine group received a weight-adjusted dose of saline, followed by free access to 4.0mL doxycycline. Forty-eight hours from the end of doxycycline supplementation, mice in the Quinine group were exposed to 1.0mL of 0.04% quinine water. Twenty-four hours later, mice were sacrificed.

##### CTA Retrieval Group

On the last day of the 4^th^ week of doxycycline supplementation, mice were trained in CTA for saccharin, as described (See Behavioral procedures), and were allowed access to 4.0mL of doxycycline-supplemented water for the rest of the day. Three days following conditioning, mice in the CTA Retrieval group were provided with 1.0mL of the CS. Twenty-four hours later, mice were sacrificed following a choice test between water and the CS.

##### LiCl Group

On the last day of doxycycline supplementation, mice received 1.0mL of 0.5% saccharin followed by an intraperitoneal injection of saline (1.5% bodyweight). For the remaining of the day, mice were allowed 4.0mL of doxycycline-water. Forty-eight hours later, mice were intraperitoneally injected with LiCl (0.14M, 1.5% body weight).

#### Extinction and Reinstatement Groups

In studies where we captured CRAM and RAM in PV-Cre mice in response to CTA Extinction and Reinstatement (Figure 7, Suppl. Fig 6 & 7), mice were provided with doxycycline supplemented water (0.5g/L) for a total of 5 weeks. On the 3^rd^ week, mice were trained to drink doxycycline from pipettes through 2 daily 1-hour sessions at the start and end of the light cycle.

On the first day of the 4^th^ week of doxycycline supplementation, mice were trained in CTA for saccharin, and were then allowed access to 4.0mL of doxycycline-supplemented water for the rest of the day. Three days following conditioning mice underwent an unreinforced choice test between the CS and water, followed by 2.0mL of doxycycline. This procedure was repeated for 9 days and behavior was recorded (extinction training).

In the last 2 days of the 5^th^ week of doxycycline supplementation, mice were allowed 4.0mL of doxycycline-water/day. Mice in both groups were then allowed 48 hours of two 1-hour water-drinking sessions/day. At the beginning of the second day of Dox clearance, mice the reinstatement group were re-exposed to an identical body weight-adjusted dose of the US, while the extinction group received saline. A final choice test between the CS and water was conducted on the next day. Behavior was recorded and mice were sacrificed 24 hours later.

### Imaging and Quantification

Images were processed using CellSens Dimension® Olympus Life Sciences® and identical regions of interest from the bilateral IC were quantified for each slice using image J® (approx. 1000um x 1000um). Representative images were additionally processed using the 2-D deconvolution® function of the CellSens software (10 iterations, 64 pixels). Slices were aligned in relation to Bregma and cell count data analyzed using GraphPad Prism®.

Slice- and Bregma aligned cell count data was further normalized across all 6 treatment groups. Normalized responses for RAM and CRAM were then analyzed separately using a 2-way ANOVA. To examine the correlation between the two responses across the IC, we compared the ratio of the normalized CRAM:RAM responses across all treatments. CRAM responses were also expressed of RAM cells/slice, and analyzed using the same framework, without further normalization. Similarly, Bregma-aligned cell count data was obtained from three PV-Cre mice (eight 35um-thick slices), previously treated to express DREADDs at the IC in a Cre-dependent manner (Suppl. Figure 2). Cell count data was scored in relation to the layers and subdivisions of the IC and plotted in relation to Bregma.

### RNAscope and Imaging

Mice were sacrificed and brains were immediately flash frozen. 20 µm sections were then made of the fresh frozen brains with a Leica CM 1950 cryostat, and mounted on SuperFrost Ultra Plus Adhesion slides (Thermo Fisher). RNAscope (ACD) in situ hybridization was then done as per the manufacturer’s protocol. Slices were fixed for 30 minutes in 4% paraformaldehyde solution and washed for 5 minutes, three times in PBS. Next, slices were placed in a series of 50, 70 and 100% ethanol washes for 5 minutes each, with a repetition of the final step at 100%. Using a hydrophobic pen, outlines were drawn around each slice and protease IV was applied for 25 min. Slides were then washed for 10-15 minutes in PBS, and were then incubated with C1-3 probes (GAD, SST, PV and VIP) for 2 h at 40°C. The slides were then washed in washing buffer for 5 min, and Amp1 was added and incubated for 30 minutes at 40°C, followed by a further 5 minutes wash in washing buffer. Amp2 was added and an incubation at 40°C was done for 15 minutes, followed by 5 minutes in washing buffer. Amp3 was then added for a 30 minutes incubation at 40°C, and another 5 minutes wash in washing buffer. Finally, we added Amp4 and incubated for 15 minutes at 40°C. Following a final wash step, slices were incubated in DAPI for 30 seconds. Slices were mounted using ProLong Gold antifade (Thermo Fisher). Images were acquired as a 3 layered z stack, 1.5 µm apart, with an Olympus IX81 microscope and CellSens software was used to deconvolute the images (Suppl. Figure 1C).

## Author Contributions

AY and KR planned the research and wrote the manuscript. KR supervised the research. AY conducted all stereotaxic surgeries and behavioral studies with the help of SKC, HK and MK. SKC carried out all electrophysiological studies. Immunohistological studies of activity-dependent transcription were carried out by AY. NG performed studies using RNA Scope. Animal welfare and maintenance by MK. All authors provided critical input to the manuscript.

## Acknowledgments

The authors would like to thank all current members of the Rosenblum lab for their help and support. Special thanks to Dr. Hanoch Kaphzan, Federica Cruciani, Gaia Auerbach and Gila Scherer for critical reading of the manuscript. This research was supported by a grant from the Israel Science Foundation (ISF); ISF 946/17; ISF 258/20; ISF-UGC 2311/15; and TransNeuro ERANET JPND supported by the Israel Ministry of Health Grant 3-14616 to K.R.

---

**Suppl. Figure 1: GABAergic interneurons exhibit distinct cell-type specific distribution across the aIC, and differentially contribute to CTA memory acquisition and retrieval**

A: Representative images from RNA scope studies in WT male mice, demonstrating differential topographical arrangement of cortical interneuronal subtypes across the layers and subregions of the aIC. B: B: Representative images from GAD-CRE mice (n=4) treated to express chemogenetic inhibitory receptors in inhibitory neurons of the aIC (aIC GAD hM4DGi). C: Representative images from SST-Cre mice (n=4) treated to express chemogenetic inhibitory receptors in excitatory neurons of the aIC (aIC SST hM4DGi). D: Representative images from VIP-CRE mice (n=4) treated to express chemogenetic inhibitory receptors in inhibitory neurons of the aIC (aIC VIP hM4DGi). E: Inhibition of aIC SST interneurons either during CTA memory acquisition (Saline – 93.284+/−1.729%, n=9; CNO – 91.012+/−1.185%, n=9) or retrieval (Saline – 91.585+/− 1.641%, n=8; CNO – 88.226+/−2.084%, n=8), did not affect CS avoidance. F: Inhibition of aIC VIP interneurons during CTA memory acquisition enhanced aversive responses during retrieval (Saline – 85.097+/−2.042, n=7; CNO – 95.687+/−2.100%, n=7). Silencing of aIC VIP during memory retrieval did not affect CS avoidance (Saline – 89.225+/−1.499, n=8; CNO – 88.164+/− 1.991%, n=10). All images acquired at Bregma 0.86. Statistical differences were assessed using 2-way ANOVA (p<0.05(*), p<0.01(**), p<0.001(***), p<0.0001(****)).

**Suppl. Figure 2: Cre-dependent cell-type specific expression of chemogenetic tools in aIC PV**

A-B: Representative images from PV-Cre mice stereotactically treated with AAVs resulting in EF1-driven, Cre-dependent expression of excitatory (hM4DGq) or inhibitory (hM4DGi) chemogenetic receptors tethered to mCherry in PV interneurons of the aIC. C-D: Quantification of aIC PV topography using 8 Bregma-aligned slices from mice treated to express hM4DGi/mCherry at the aIC (n=3). C: The majority of aIC PV interneurons are located within the granular insular cortex (GI – 63.29+/−3.010cells/slice). A smaller but stable number of aIC PV was localized within the dysgranular (DI – 45.25+/−1.900cells/slice) and agranular (AI – 38.83+/− 1.127cells/slice) sub-regions. D: By comparing the layer-specific localization of aIC PV we show that PV primarily reside within deep- (LV-VI – 116.2+/−3.971cells/slice), and to a lesser extent in superficial layers of the aIC (LI-III – 30.96+/−2.408cells/slice).

**Suppl. Figure 3: Appetitive taste memory retrieval increases activity-dependent transcription at the aIC compared to aversive tastants**

A: The CRAM/RAM TET OFF system was utilized in order to label activity-dependent transcription in the aIC of PV-Cre mice (see Methods). Activity-dependent transcription was captured in mice following Saccharin Retrieval (n=4), Quinine (n=4), as well as CTA Retrieval (n=5). B-D: Representative images from the three treatment groups. Channels displayed from left to right: DAPI, EGFP RAM, Merge. Scale bars represent 100um. E: Bilateral RAM cell count data from 8 consecutives slices from the three treatments was plotted in relation to Bregma. Mean RAM response across the aIC were significantly increased in the Saccharin Retrieval group (155.0+/−7.977cells/slice) compared to both Quinine (115.3+/−2.219cells/slice) and CTA Retrieval (122.6+/−3.239cells/slice). F: Normalized mean RAM responses among the three treatments. G: Behavioral assessment of saccharin aversion in the Saccharin (31.37+/−7.203%, n=4) and CTA retrieval (83.83+/−4.385%, n=5) groups. Statistical differences were assessed using 2-way ANOVA (E-F) and Unpaired t-test (G), respectively (p<0.05(*), p<0.01(**), p<0.001(***), p<0.0001(****)).

**Suppl. Figure 4: Aversive taste memory retrieval increases the recruitment of aIC PV in the activated ensemble**

A: The CRAM/RAM TET OFF system was utilized in order to label activity-dependent transcription in aIC PV interneurons of PV-Cre mice (see Methods). Activity-dependent transcription was captured in mice following appetitive taste memory retrieval (Saccharin Retrieval, n=4), aversive taste memory retrieval (CTA retrieval, n=5) as well as in response to innately aversive taste exposure (Quinine, n=4). B: Bilateral CRAM cell count data from 8 consecutives slices from the three treatments was expressed as a percentage of the RAM response, and plotted in relation to Bregma (%CRAM/RAM). C: We examined the correlation between CRAM and RAM responses across the treatment groups, from 8 consecutive slices and in relation to Bregma. D: Mean %CRAM/RAM response across the aIC was significantly increased in response to CTA Retrieval (37.89+/−1.142%) compared to both Quinine (16.02+/−1.713%), and Saccharin Retrieval (3.613+/−0.4818%). Mean %CRAM/RAM responses in the Quinine group were also elevated compared to Saccharin Retrieval. E: Mean correlation between CRAM and RAM was highest in the CTA retrieval group (1.099+/−0.03312) compared to both Quinine (0.4648+/−0.04968) and Saccharin Retrieval (0.1273+/−0.02367). The correlation between CRAM and RAM in the Quinine was also significantly different to Saccharin Retrieval. Statistical differences between the groups were assessed using 2-way ANOVA (p<0.05(*), p<0.01(**), p<0.001(***), p<0.0001(****)).

**Suppl. Figure 5: PV-dependent modulation of mIPSCs on the aIC-BLA projection is necessary for CTA reinstatement following extinction**

A: Schematic of behavioral proceedings for mice undergoing Extinction and Reinstatement prior to electrophysiological recordings. Representative traces of mIPSCs from mice following Extinction and Reinstatement. Scale Bar represents 2sec and 50pA. B: Schematic of behavioral proceedings and graphical plot of aversion during Extinction and Reinstatement in WT mice (n=8/group). Aversion across the two groups was similar during CTA retrieval (Extinction – 88.813+/−2.487%; Reinstatement – 91.288+/−1.622%), and following Extinction (Extinction – 25.063+/−2.251%; Reinstatement – 24.475+/−2.111%). Reinstatement significantly increased aversion compared to Extinction during a subsequent memory test (Extinction – 34.824+/−4.824%; Reinstatement – 78.2053+/−2.053%).

**Suppl. Figure 6: Extinction induces activity-dependent transcription at the aIC compared to reinstatement**

A: The CRAM/RAM TET OFF system was utilized in order to label activity-dependent transcription in the aIC of PV-Cre mice (see Methods). Activity-dependent transcription was captured in mice following CTA Extinction (n=3), Reinstatement (n=3), as well as LiCl (n=4). B-D: Representative images from the three treatment groups. Channels displayed from left to right: DAPI, EGFP RAM, Merge. Scale bars represent 100um. E: Bilateral RAM cell count data from 8 consecutives slices from the three treatments was plotted in relation to Bregma. Mean RAM response across the aIC were significantly increased in the Extinction group (181.2+/− 6.088cells/slice) compared to both Reinstatement (141.2+/−6.081cells/slice) and LiCl (70.88+/− 3.513cells/slice). F: Normalized mean RAM responses among the three treatments. G: Behavioral assessment of saccharin aversion in the Extinction (45.573+/−4.854%, n=3) and Reinstatement (84.642+/−3.937%, n=3) groups. Statistical differences were assessed using 2-way ANOVA (E-F) and Unpaired t-test (G), respectively (p<0.05(*), p<0.01(**), p<0.001(***), p<0.0001(****)).

**Suppl. Figure 7: Reinstatement increases the recruitment of aIC PV interneurons in the activated ensemble**

A: The CRAM/RAM TET OFF system was utilized in order to label activity-dependent transcription in aIC PV interneurons of PV-Cre mice (see Methods). Activity-dependent transcription was captured in mice following CTA Extinction (n=4), Reinstatement (n=5), as well as in response to LiCl exposure alone (n=4). B: Bilateral CRAM cell count data from 8 consecutives slices from the three treatments was expressed as a percentage of the RAM response, and plotted in relation to Bregma (%CRAM/RAM). C: We examined the correlation between CRAM and RAM responses across the treatment groups, from 8 consecutive slices and in relation Bregma. D: Mean %CRAM/RAM response across the aIC was significantly increased in response to Reinstatement (31.79+/−2.937%) and LiCl (24.49+/−1.161%) groups was increased compared to Extinction (9.187+/−1.226%). E: Mean correlation between CRAM and RAM was increased in the Reinstatement group (1.099+/−0.03312) and LiCl (0.1273+/−0.02367) groups compared to Extinction (0.4648+/−0.04968). Statistical differences between the groups were assessed using 2-way ANOVA (p<0.05(*), p<0.01(**), p<0.001(***), p<0.0001(****)).

| TABLE OF STATISTICS: MAIN FIGURES |  |  |
| --- | --- | --- |
| Figure | Statistical Test | Results |
| 1C | 2-way ANOVA<br>Male aIC CaMKII Inhibition vs. Controls (+/- CNO)<br>Sidak's Multiple Comparisons<br>Acquisition and Retrieval | Treatment; $p=0.0002$, $F(1, 32) = 17.37$<br>Timing; $p=0.0974$, $F(1, 32) = 2.917$<br>Interaction; $p=0.4046$, $F(1, 32) = 0.7133$<br>Acquisition: $p=0.0496$, $t=2.350$, $df=32$<br>Retrieval: $p=0.0025$, $t=3.545$, $df=32$ |
| 1D | 2-way ANOVA<br>Male aIC GAD Inhibition vs. Controls (+/- CNO)<br>Sidak's Multiple Comparisons<br>Acquisition and Retrieval | Treatment; $p<0.0001$, $F(1, 41) = 19.47$<br>Timing; $p<0.0001$, $F(1, 41) = 40.65$<br>Interaction; $p<0.0001$, $F(1, 41) = 68.72$<br>Acquisition: $p=0.0232$, $t=2.642$, $df=41$<br>Retrieval: $p<0.0001$, $t=9.349$, $df=41$ |
| 1E | 2-way ANOVA<br>Male Saline Controls vs. CNO Controls (+/- CNO)<br>Sidak's Multiple Comparisons<br>Acquisition and Retrieval | Treatment; $p=0.8159$, $F(1, 40) = 0.05490$<br>Timing; $p=0.5993$, $F(1, 40) = 0.2805$<br>Interaction; $p=0.2551$, $F(1, 40) = 1.333$<br>Acquisition: $p=0.7686$, $t=0.6507$, $df=40$<br>Retrieval: $p=0.5537$, $t=0.9821$, $df=40$ |
| 2C | 2-way ANOVA<br>Amplitude<br>Sidak's Multiple Comparisons<br>CTA Retrieval vs. Saccharin Retrieval<br>aIC neurons and aIC-BLA projecting neurons | Treatment; $p=0.0054$, $F(1, 47) = 8.506$<br>Population; $p=0.3825$, $F(1, 47) = 0.7771$<br>Interaction; $p=0.4116$, $F(1, 47) = 0.6864$<br>aIC neurons: $p=0.2901$, $t=1.437$, $df=47$<br>aIC-BLA projecting neurons: $p=0.0179$, $t=2.726$, $df=47$ |
| 2D | 2-way ANOVA<br>Frequency<br>Sidak's Multiple Comparisons<br>CTA Retrieval vs. Saccharin Retrieval<br>aIC neurons and aIC-BLA projecting neurons | Treatment; $p<0.0001$, $F(1, 47) = 27.65$<br>Population; $p=0.3293$, $F(1, 47) = 0.9718$<br>Interaction; $p=0.01446$, $F(1, 47) = 2.201$<br>aIC neurons: $p<0.0001$, $t=4.639$, $df=47$<br>aIC-BLA projecting neurons: $p=0.0169$, $t=2.748$, $df=47$ |
| 3C | 2-way ANOVA<br>Amplitude<br>Sidak's Multiple Comparisons<br>CTA Retrieval vs. Saccharin Retrieval vs. CTA Retrieval + aIC PV Inhibition<br>aIC neurons and aIC-BLA projecting neurons | Treatment; $p=0.0205$, $F(2, 76) = 4.093$<br>Population; $p=0.9299$, $F(2, 76) = 0.007789$<br>Interaction; $p=0.0193$, $F(2, 76) = 4.157$<br>aIC neurons:<br>Saccharin Retrieval vs. CTA Retrieval<br>$p=0.5555$, $t=1.192$, $df=76$<br>Saccharin Retrieval vs. CTA Retrieval + CNO<br>$p=0.1305$, $t=2.033$, $df=76$<br>CTA Retrieval vs. CTA Retrieval + CNO<br>$p=0.7647$, $t=0.8782$, $df=76$<br>aIC-BLA projecting neurons:<br>Saccharin Retrieval vs. CTA Retrieval<br>$p=0.0136$, $t=2.923$, $df=76$<br>Saccharin Retrieval vs. CTA Retrieval + CNO<br>$p=0.9936$, $t=0.2358$, $df=76$<br>CTA Retrieval vs. CTA Retrieval + CNO<br>$p=0.0041$, $t=3.323$, $df=76$ |
| 3D | 2-way ANOVA<br>Frequency<br>CTA Retrieval vs. Saccharin Retrieval vs. CTA Retrieval + aIC PV Inhibition<br>aIC neurons and aIC-BLA projecting neurons | Treatment; $p<0.0001$, $F(2, 76) = 14.63$<br>Population; $p=0.7069$, $F(2, 76) = 0.1424$<br>Interaction; $p=0.4925$, $F(2, 76) = 0.7148$<br>aIC neurons:<br>Saccharin Retrieval vs. CTA Retrieval<br>$p=0.1854$, $t=1.865$, $df=76$<br>Saccharin Retrieval vs. CTA Retrieval + CNO<br>$p=0.7674$, $t=0.8737$, $df=76$<br>CTA Retrieval vs. CTA Retrieval + CNO<br>$p=0.0163$, $t=2.860$, $df=76$<br>aIC-BLA projecting neurons:<br>Saccharin Retrieval vs. CTA Retrieval<br>$p=0.0009$, $t=3.776$, $df=76$<br>Saccharin Retrieval vs. CTA Retrieval + CNO<br>$p=0.9641$, $t=0.4275$, $df=76$<br>CTA Retrieval vs. CTA Retrieval + CNO<br>$p<0.00001$, $t=4.430$, $df=76$ |
| 4E | 2-way ANOVA<br>Male aIC PV Inhibition vs. Controls (+/- CNO)<br>Sidak's Multiple Comparisons<br>Acquisition and Retrieval | Treatment; $p<0.0001$, $F(1, 38) = 37.72$<br>Timing; $p<0.0001$, $F(1, 38) = 27.12$<br>Interaction; $p<0.0001$, $F(1, 38) = 26.63$<br>Acquisition: $p=0.7012$, $t=0.7575$, $df=38$ |
| | | Retrieval; $p < 0.0001$, $t = 10.29$, $df = 38$ |
| 4F | 2-way ANOVA<br>CTA Retrieval vs. CTA Retrieval + aIC PV Inhibition<br>Sidak's Multiple Comparisons<br>Male and Female mice | Treatment; $p < 0.0001$, $F(1, 32) = 74.44$<br>Sex; $p = 4542$, $F(1, 32) = 0.5740$<br>Interaction; $p = 0.0748$, $F(1, 32) = 3.392$<br>Female: $p < 0.0001$, $t = 6.600$, $df = 32$<br>Male: $p < 0.0001$, $t = 5.571$, $df = 32$ |
| 4H | 2-way ANOVA<br>Sidak's Multiple Comparisons<br>CTA Extinction vs. aIC PV Inhibition During Retrieval followed by Extinction | Treatment; $p = 0.4885$, $F(1, 189) = 0.04818$<br>Time/Extinction; $p < 0.0001$, $F(8, 189) = 13.17$<br>Interaction; $p = 0.0966$, $F(8, 189) = 1.718$<br>Day 1; $p = 0.0065$, $t = 3.436$, $df = 189$<br>Day 2; $p > 0.9999$, $t = 0.4042$, $df = 189$<br>Day 3; $p > 0.9999$, $t = 0.1594$, $df = 189$<br>Day 4; $p > 0.9999$, $t = 0.2910$, $df = 189$<br>Day 5; $p > 0.9999$, $t = 0.02193$, $df = 189$<br>Day 6; $p > 0.9999$, $t = 0.3741$, $df = 189$<br>Day 7; $p > 0.9999$, $t = 0.1440$, $df = 189$<br>Day 8; $p = 0.9726$, $t = 0.9778$, $df = 189$<br>Day 9; $p = 0.9655$, $t = 1.013$, $df = 189$ |
| 5E-F & 7E-F | 2-way ANOVA<br>Normalized CRAM<br>Sidak's Multiple Comparisons<br>Saccharin Retrieval<br>Quinine<br>CTA Retrieval<br>Extinction<br>Reinstatement<br>LiCl<br>(8 consecutive slices/subject) | Treatment; $p < 0.0001$, $F(5, 136) = 40.21$<br>Bregma; $p = 0.5009$, $F(7, 136) = 0.9099$<br>Interaction; $p = 0.9962$, $F(35, 136) = 0.4517$<br>Saccharin Retrieval vs. CTA Retrieval; $p < 0.0001$, $t = 15.86$, $df = 136$<br>Saccharin Retrieval vs. Quinine; $p = 0.0102$, $t = 4.849$, $df = 136$<br>Saccharin Retrieval vs. Extinction; $p = 0.0272$, $t = 4.396$, $df = 136$<br>Saccharin Retrieval vs. Reinstatement; $p < 0.0001$, $t = 14.74$, $df = 136$<br>Saccharin Retrieval vs. LiCl; $p = 0.0140$, $t = 4.709$, $df = 136$<br>CTA Retrieval vs. Quinine; $p < 0.0001$, $t = 10.75$, $df = 136$<br>CTA Retrieval vs. Extinction; $p < 0.0001$, $t = 9.971$, $df = 136$<br>CTA Retrieval vs. Reinstatement; $p = 0.9909$, $t = 0.8488$, $df = 136$<br>CTA Retrieval vs. LiCl; $p < 0.0001$, $t = 10.90$, $df = 136$<br>Quinine vs. Extinction; $p > 0.9999$, $t = 0.09353$, $df = 136$<br>Quinine vs. Reinstatement; $p < 0.0001$, $t = 10.25$, $df = 136$<br>Quinine vs. LiCl; $p > 0.9999$, $t = 0.1399$, $df = 136$<br>Extinction vs. Reinstatement; $p < 0.0001$, $t = 9.678$, $df = 136$<br>Extinction vs. LiCl; $p > 0.9999$, $t = 0.03597$, $df = 136$<br>Reinstatement vs. LiCl; $p < 0.0001$, $t = 10.38$, $df = 136$ |
| 6B | Unpaired t-test<br>Amplitude aIC-BLA projecting neurons<br>Extinction vs. Reinstatement | $p = 0.2135$, $t = 1.281$, $df = 22$ |
| 6C | Unpaired t-test<br>Frequency aIC-BLA projecting neurons<br>Extinction vs. Reinstatement | $p = 0.0095$, $t = 2.828$, $df = 23$ |
| 6D-E | 2-way ANOVA<br>Control RI vs. Control Extinction vs. Reinstatement + aIC PV Inhibition vs. Extinction + aIC PV Activation<br>Sidak's Multiple Comparisons<br>CTA, Extinction and Test | Treatment; $p < 0.0001$, $F(3, 108) = 27.17$<br>Test Session; $p < 0.0001$, $F(2, 108) = 199.8$<br>Interaction; $p < 0.0001$, $F(6, 108) = 16.64$<br>CTA:<br>Reinstatement + aIC PV Inhibition vs. Control Reinstatement<br>$p = 0.7698$, $t = 1.242$, $df = 108$<br>Reinstatement + aIC PV Inhibition vs. Extinction + aIC PV Activation<br>$p = 0.9307$, $t = 0.9210$, $df = 108$<br>Reinstatement + aIC PV Inhibition vs. Control Extinction<br>$p = 0.9995$, $t = 0.3641$, $df = 108$<br>Control Reinstatement vs. Extinction + aIC PV Activation<br>$p = 0.9985$, $t = 0.4390$, $df = 108$<br>Control Reinstatement vs. Control Extinction<br>$p = 0.4230$, $t = 1.724$, $df = 108$<br>Extinction + aIC PV Activation vs. Control Extinction<br>$p = 0.6326$, $t = 1.437$, $df = 108$<br>Extinction:<br>Reinstatement + aIC PV Inhibition vs. Control Reinstatement<br>$p = 0.9988$, $t = 0.4245$, $df = 108$<br>Reinstatement + aIC PV Inhibition vs. Extinction + aIC PV Activation<br>$p = 0.3416$, $t = 1.848$, $df = 108$<br>Reinstatement + aIC PV Inhibition vs. Control Extinction<br>$p = 0.6103$, $t = 1.467$, $df = 108$<br>Control Reinstatement vs. Extinction + aIC PV Activation<br>$p = 0.6717$, $t = 1.383$, $df = 108$<br>Control Reinstatement vs. Control Extinction<br>$p = 0.9000$, $t = 1.002$, $df = 108$<br>Extinction + aIC PV Activation vs. Control Extinction |
|  |  | p=0.9987, t=0.4267, df=108<br>Test:<br>Reinstatement + aIC PV Inhibition vs. Control Reinstatement<br>p<0.0001, t=7.880, df=108<br>Reinstatement + aIC PV Inhibition vs. Extinction + aIC PV Activation<br>Reinstatement + aIC PV Inhibition vs. Control Extinction<br>p<0.0001, t=8.484, df=108<br>Control Reinstatement vs. Extinction + aIC PV Activation<br>p=0.9340, t=0.9111, df=108<br>Control Reinstatement vs. Control Extinction<br>p>0.9999, t=0.1487, df=108<br>Extinction + aIC PV Activation vs. Control Extinction<br>p<0.0001, t=10.50, df=108 |

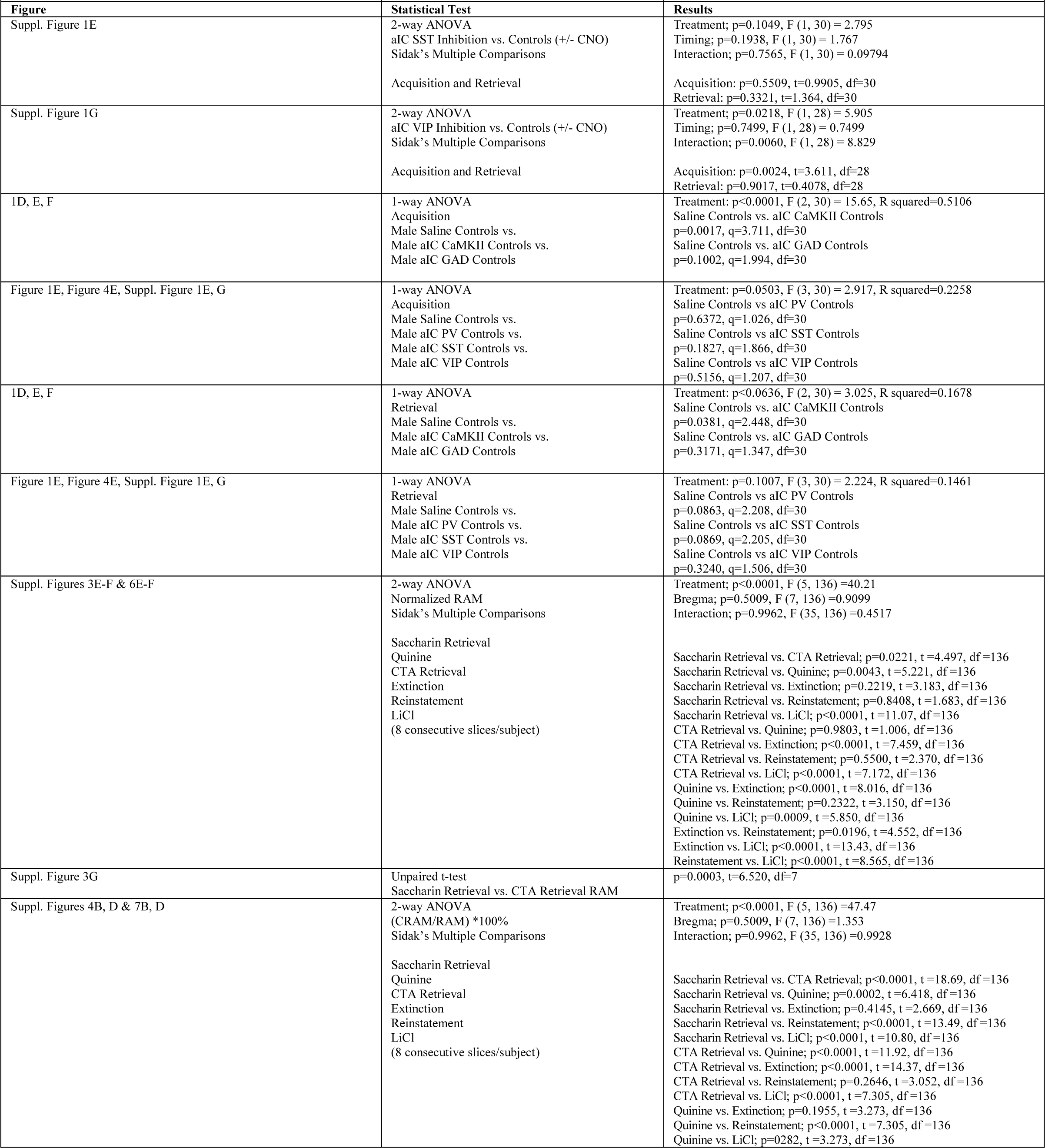
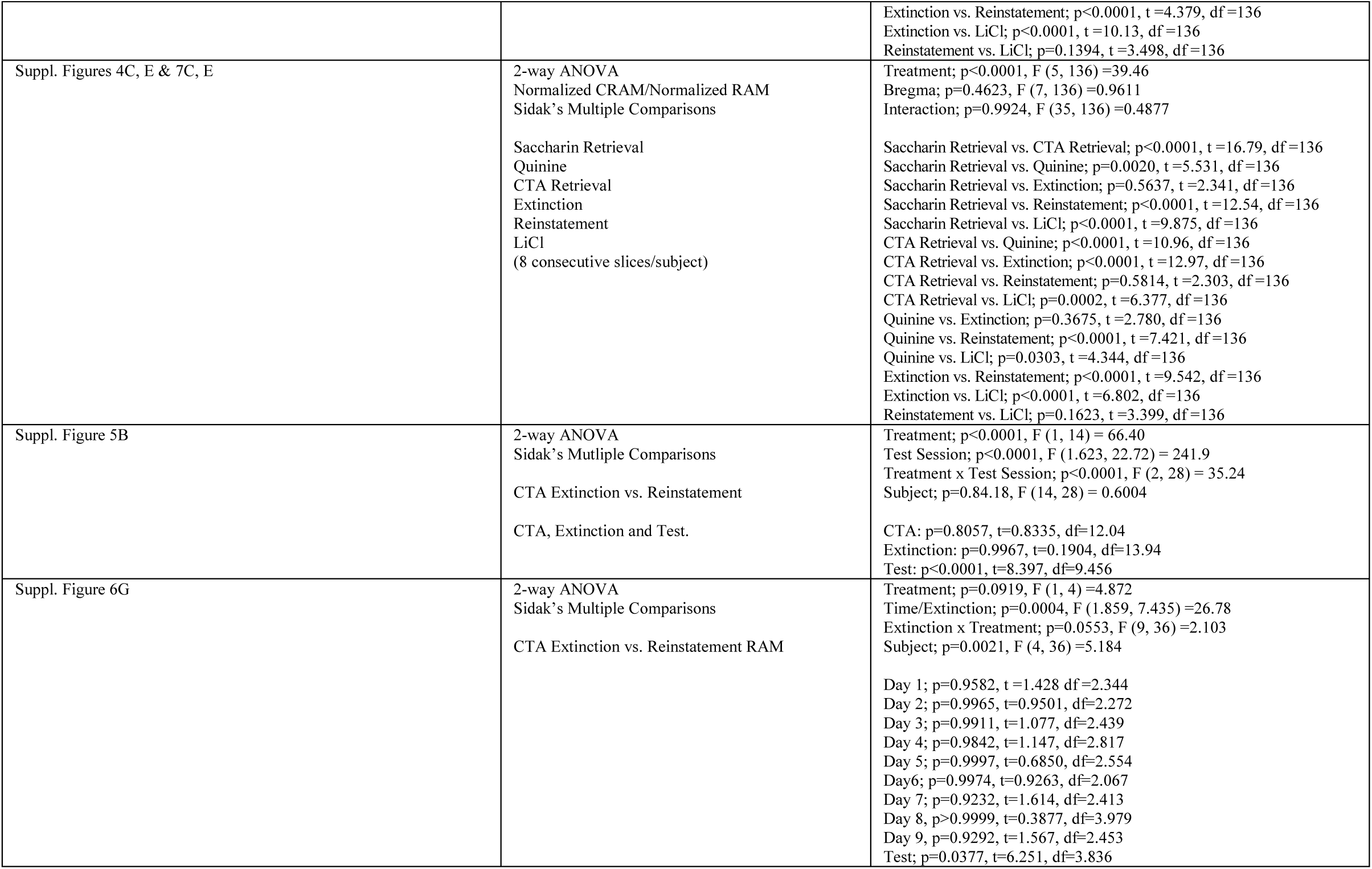

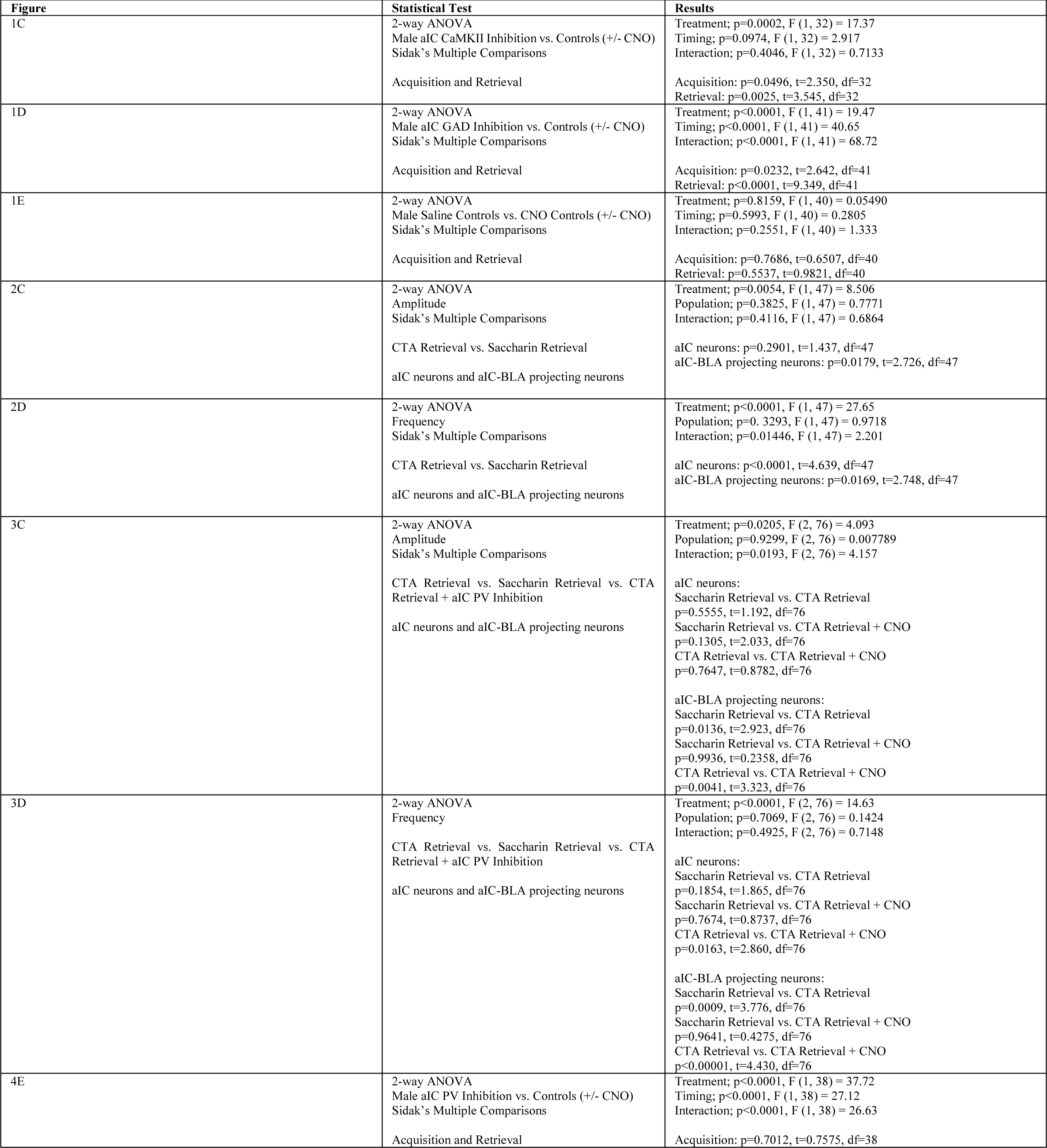
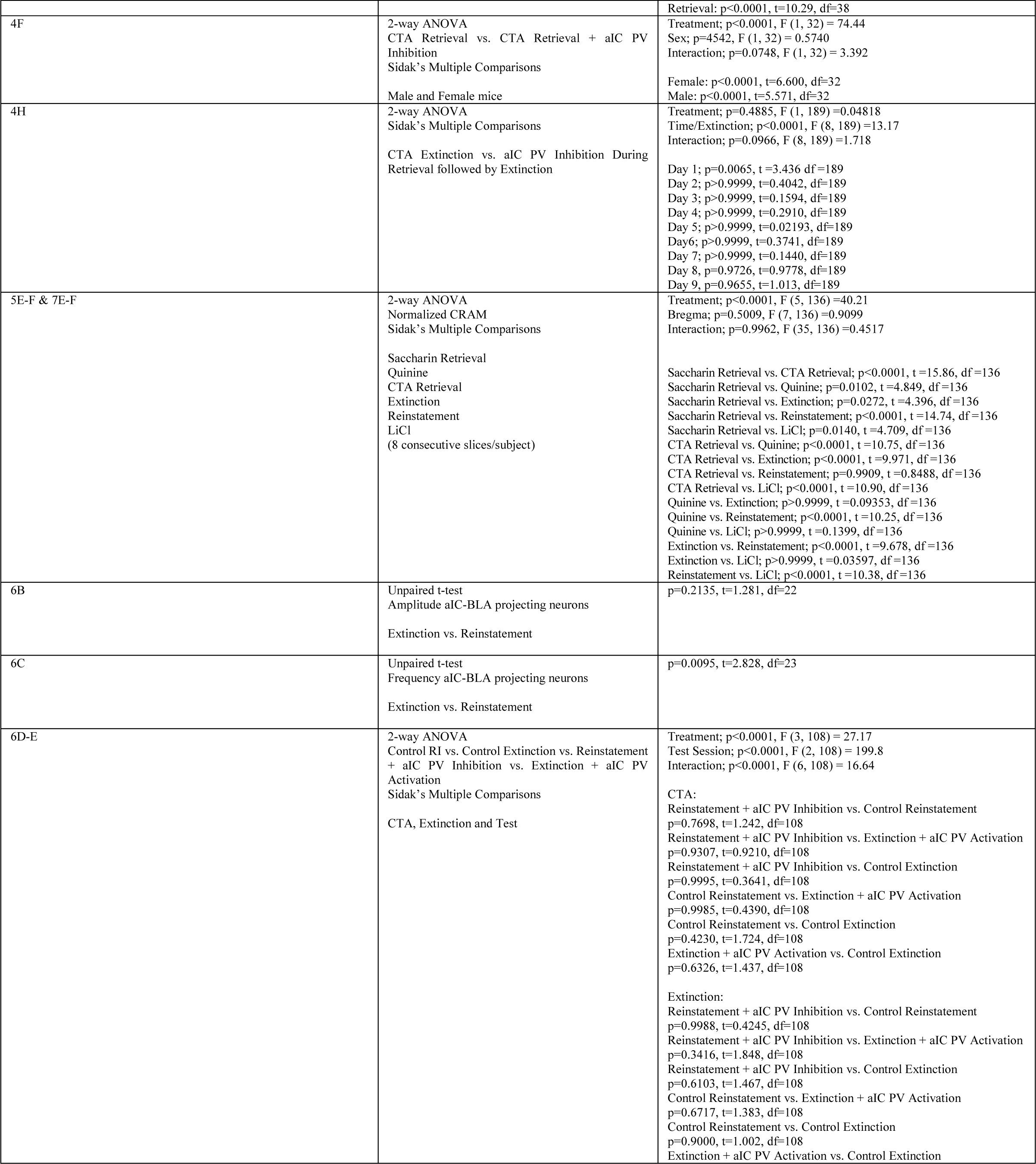
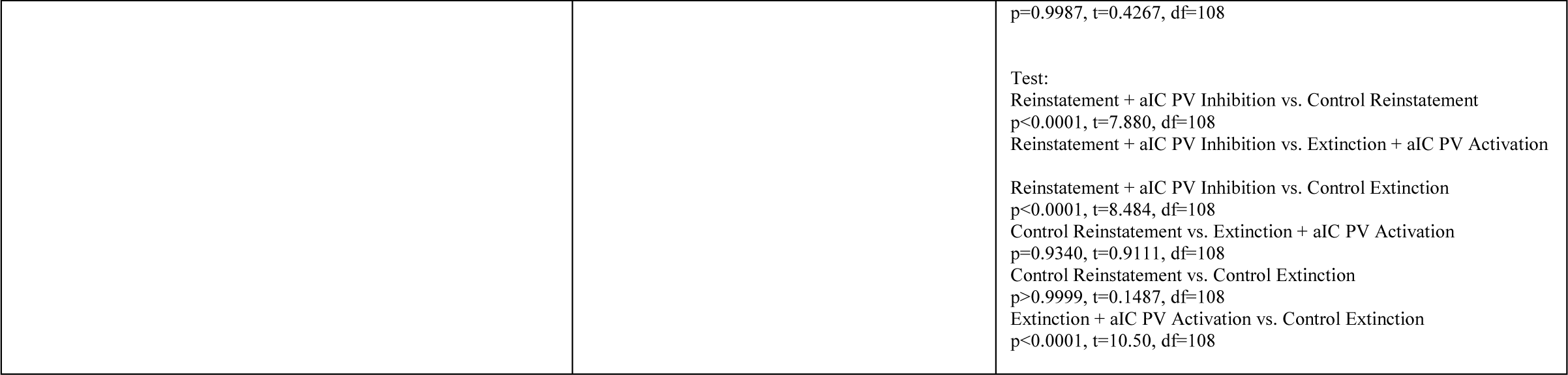

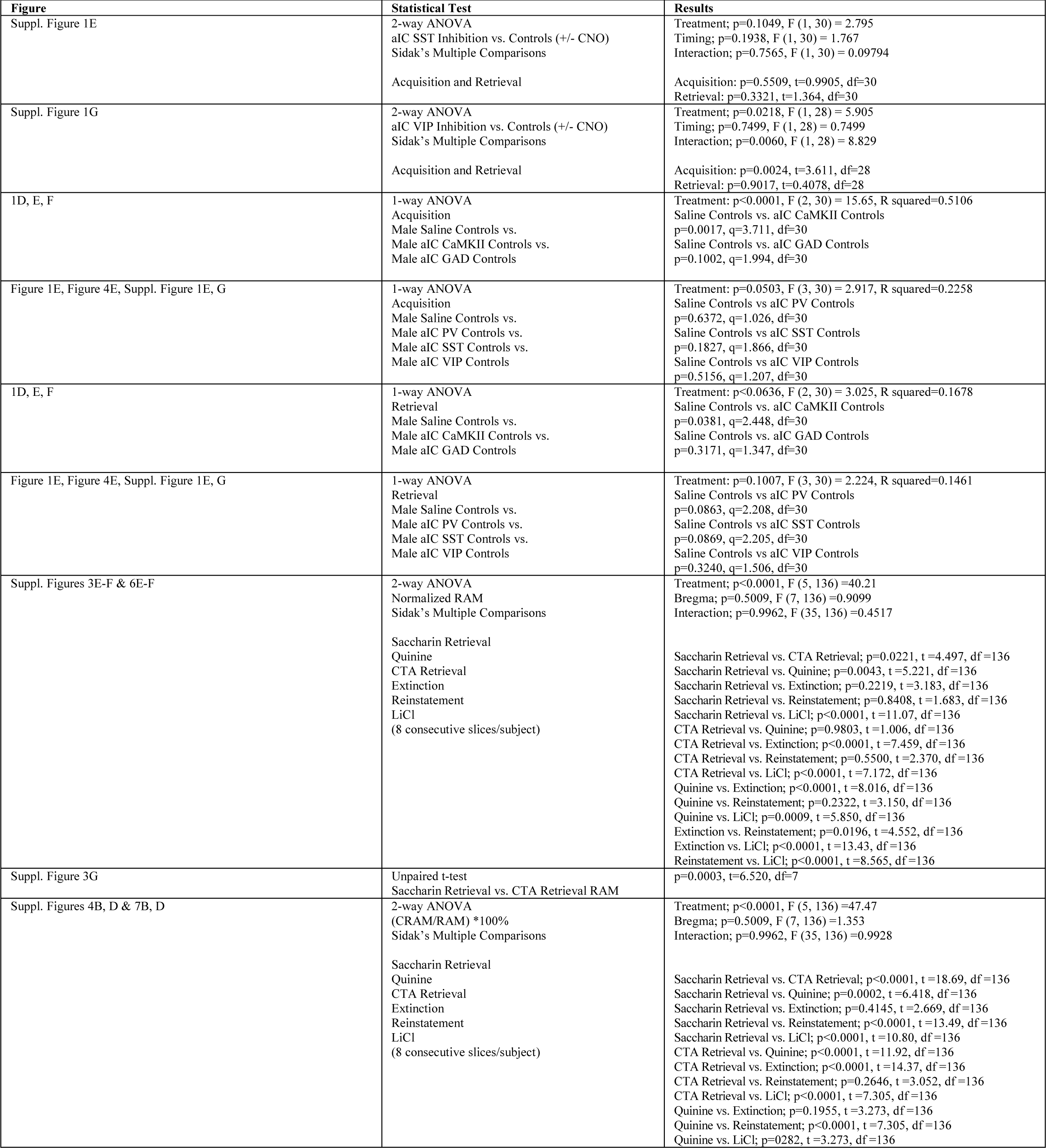
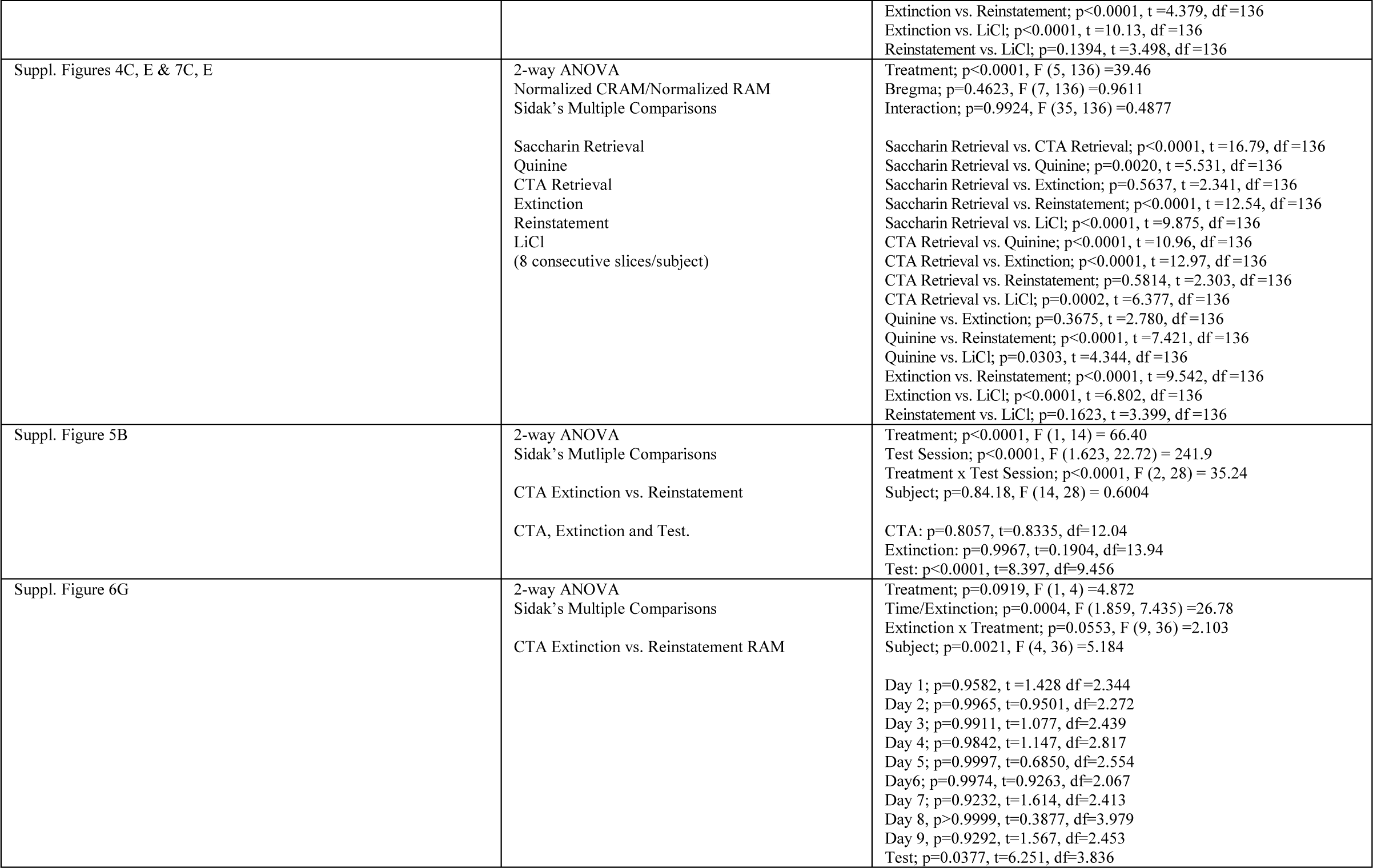

