## Supplementary figures and images for "Parvalbumin interneuron inhibition onto anterior insula neurons projecting to the basolateral amygdala orchestrates aversive taste memory retrieval"

### supplementary figures

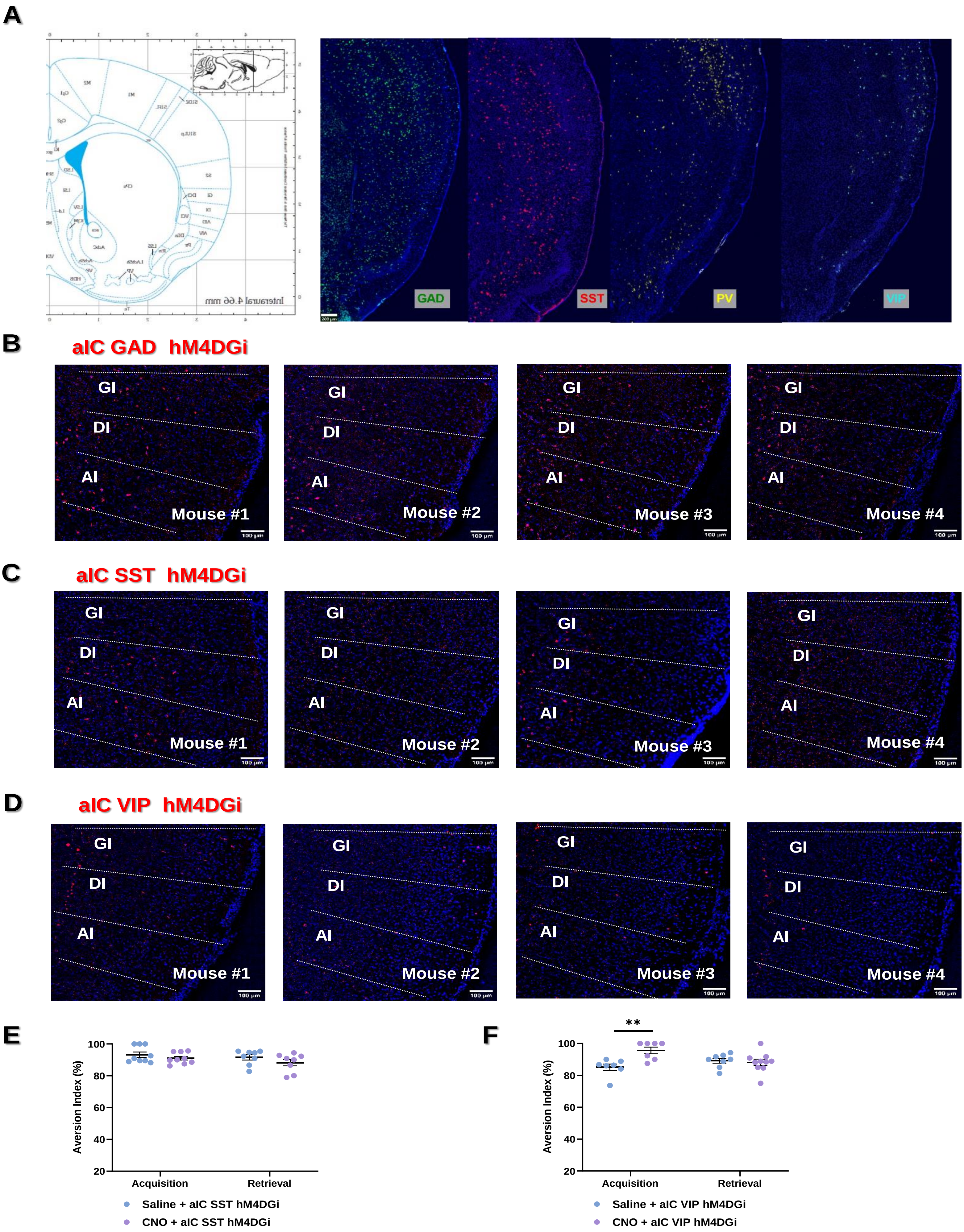

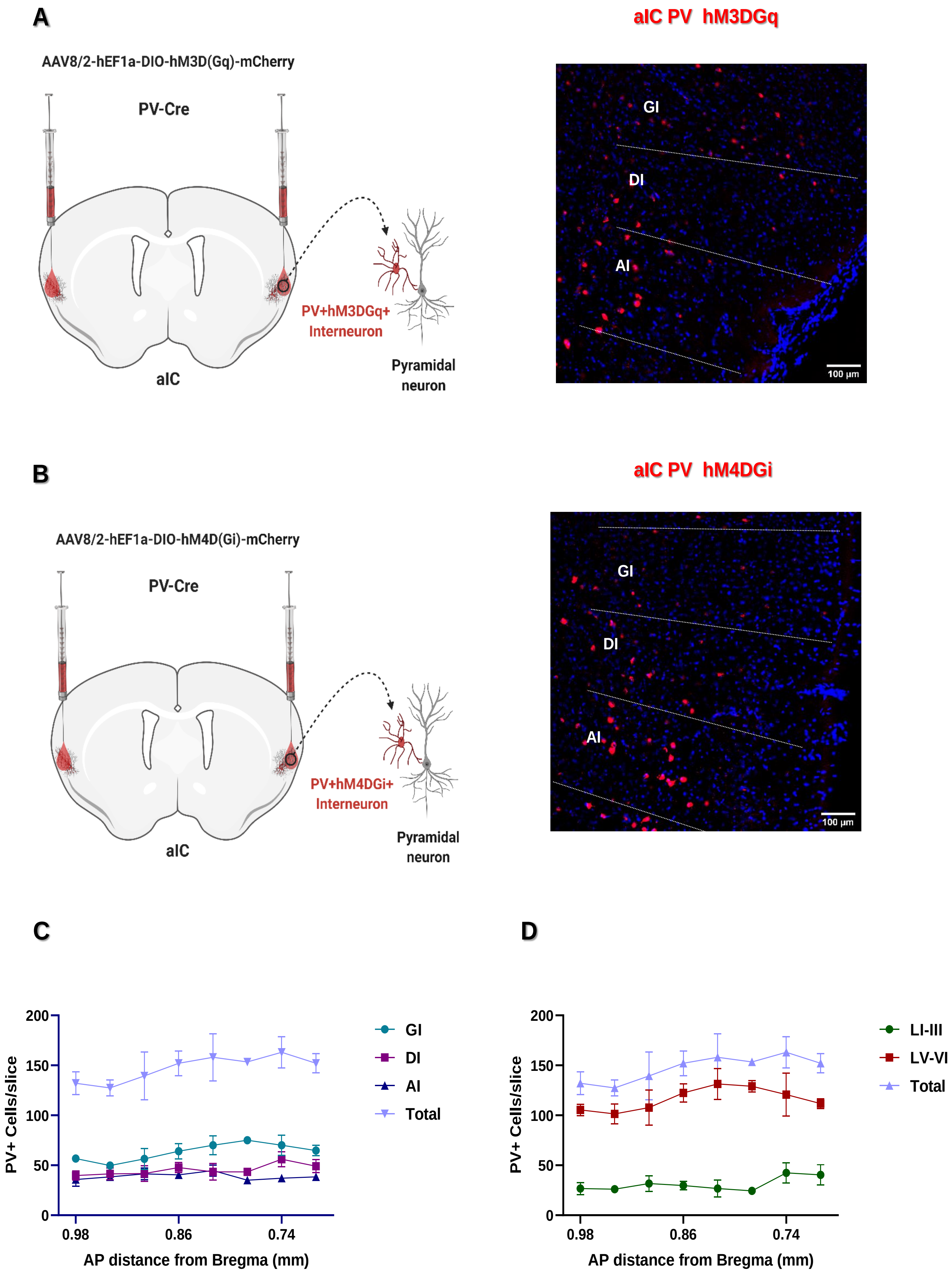

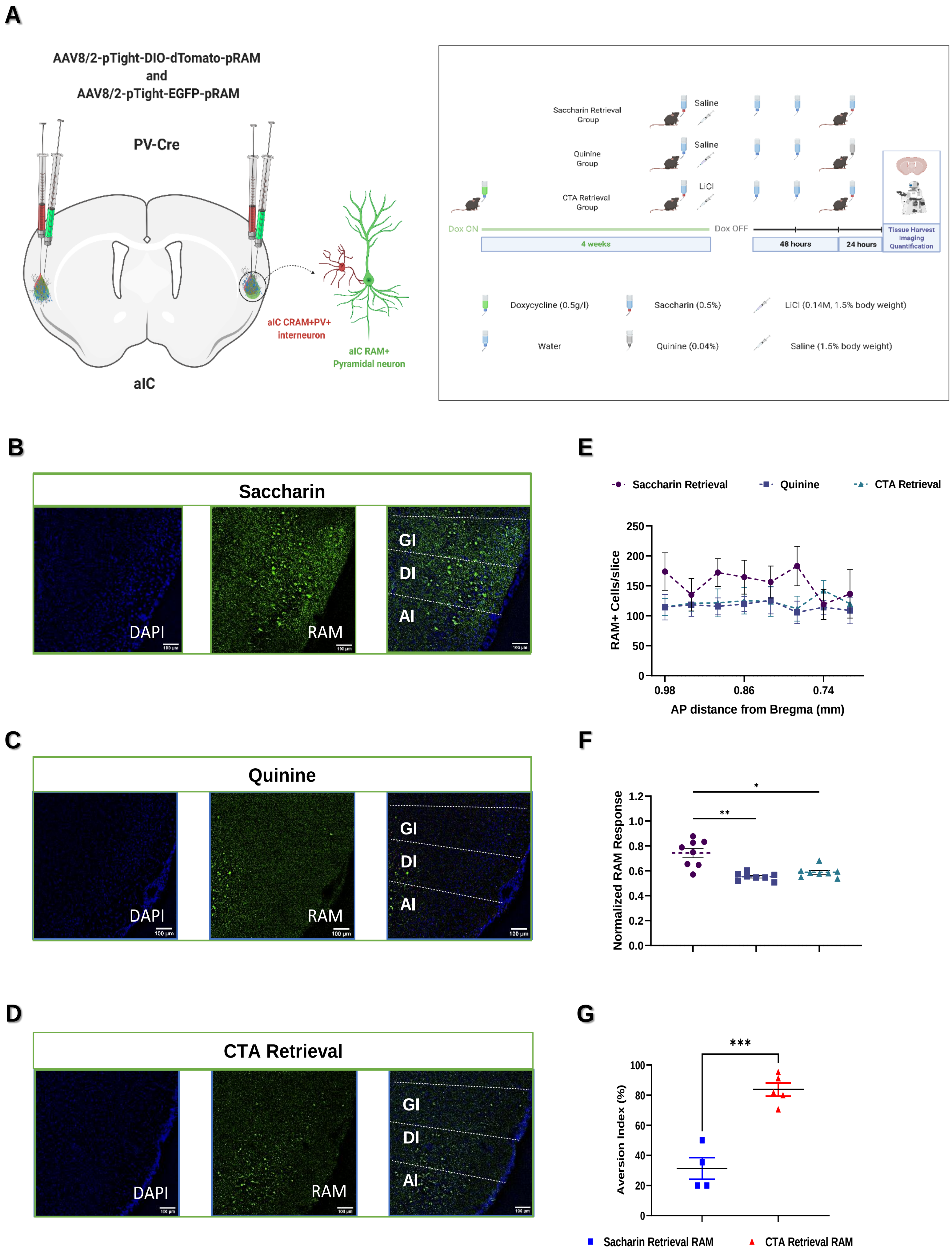

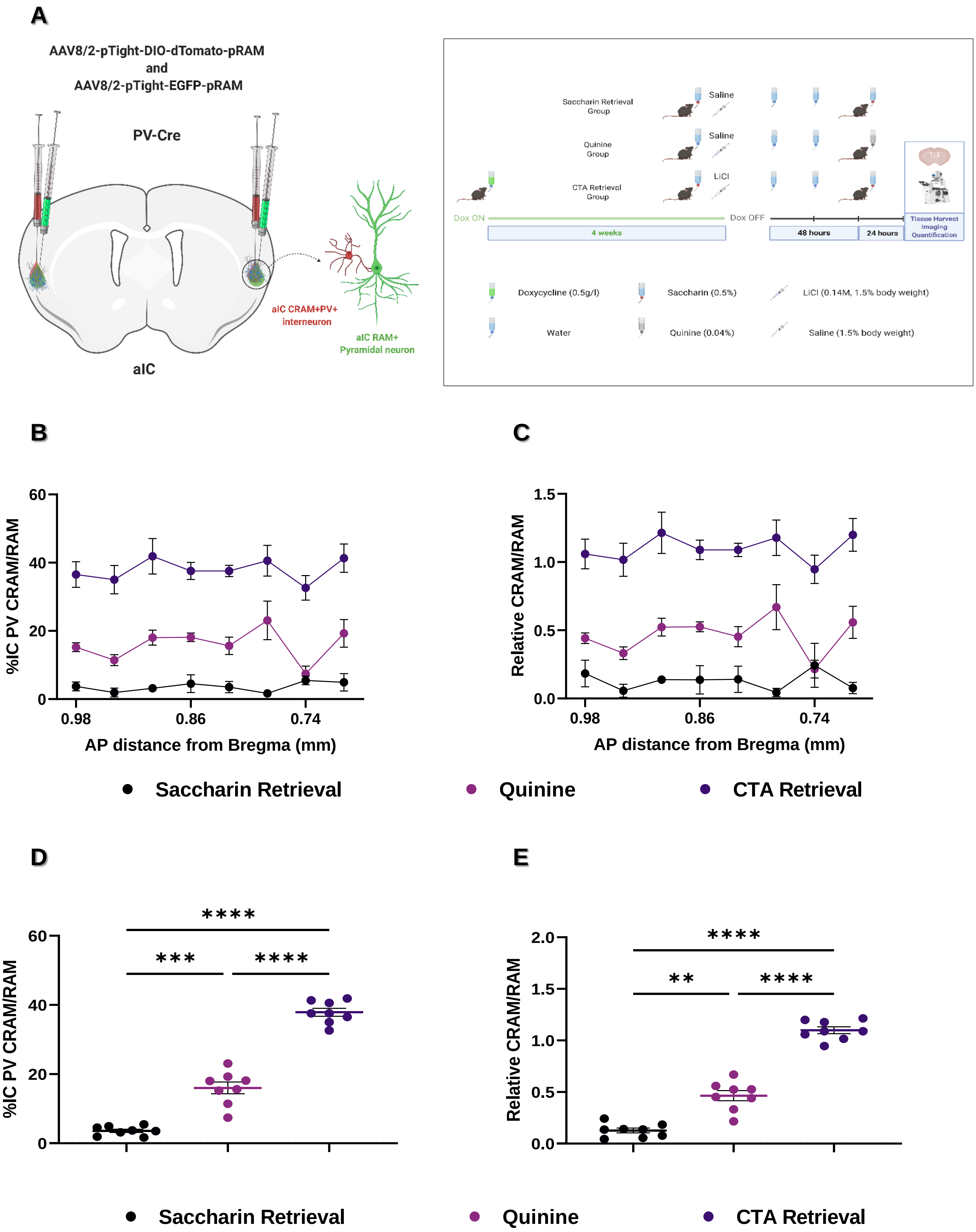

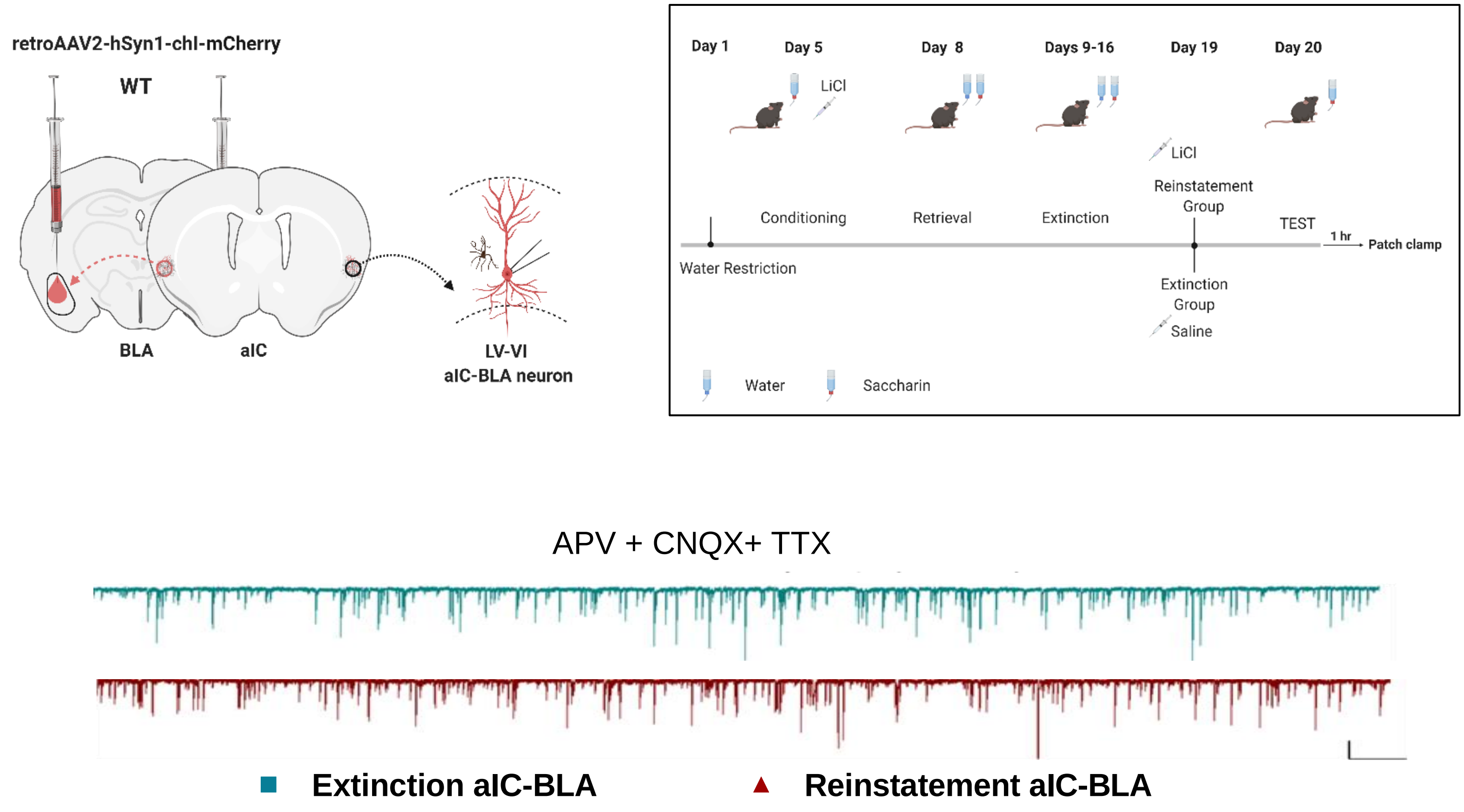

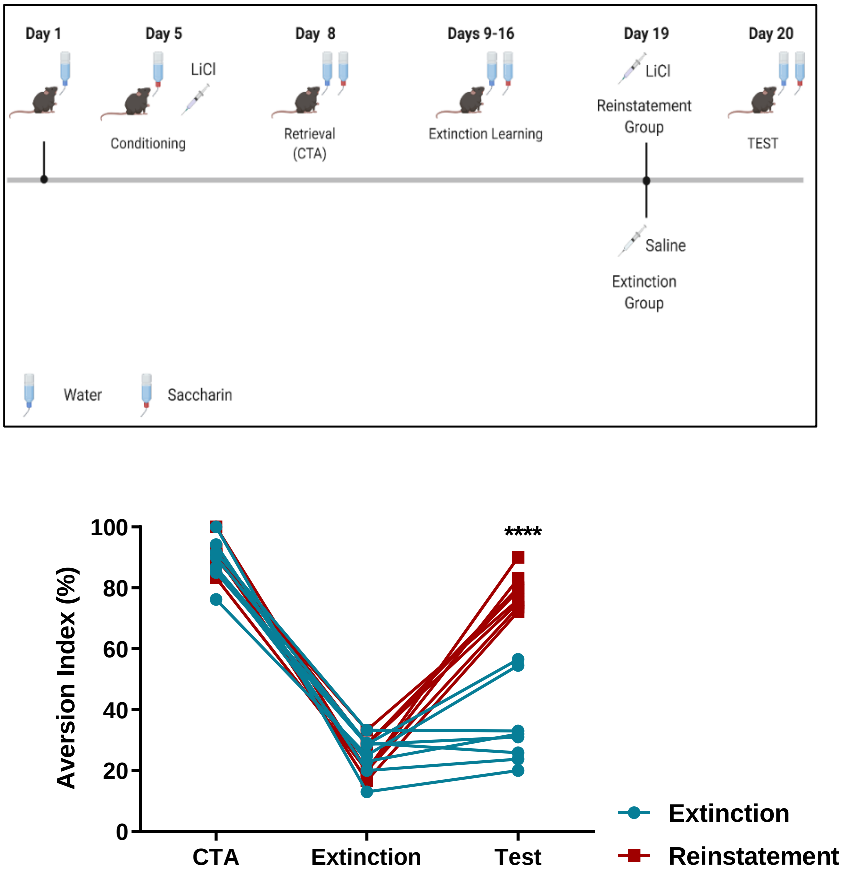

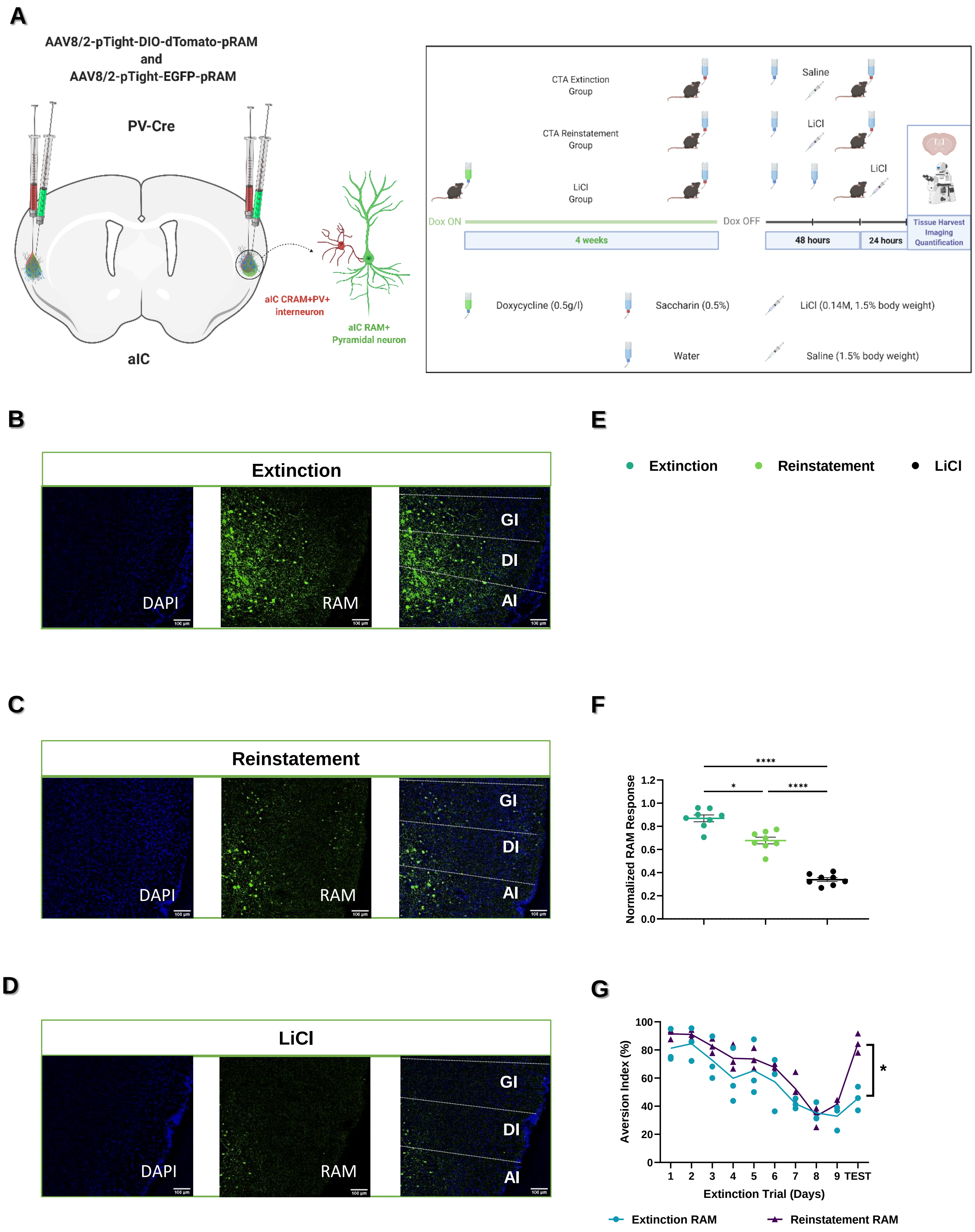

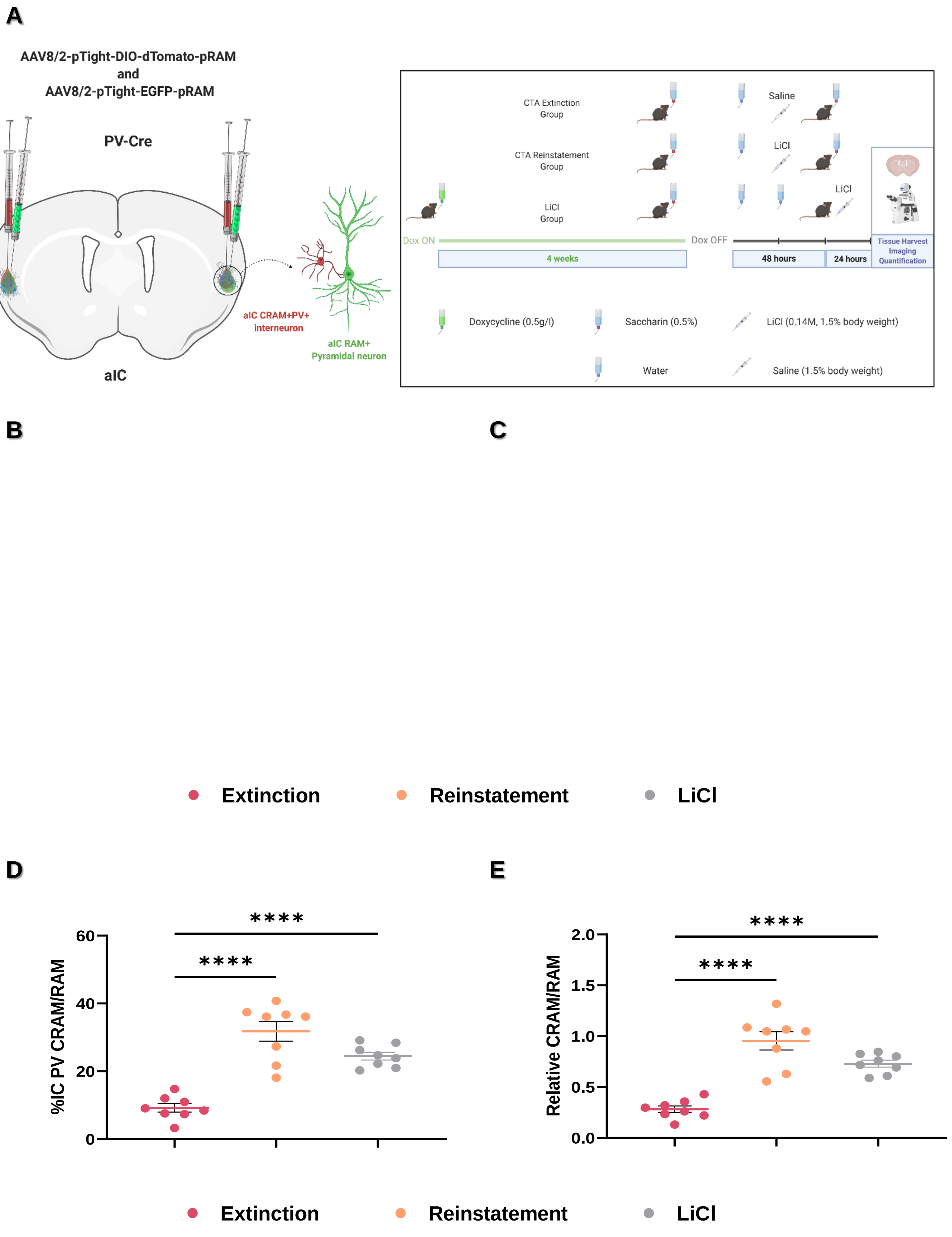
